# Weak pervasive incompatibilities and compensatory adaptation drive hybrid genome evolution in yeast

**DOI:** 10.64898/2026.01.07.698244

**Authors:** Artemiza A. Martínez, Gregory I. Lang

## Abstract

Species barriers limit gene flow and maintain co-adapted genomes. Interspecific hybridization can break down species barriers to reveal genetic incompatibilities. Although the phenotypic and genomic consequences of hybridization have been extensively studied in many systems, far less is known about the longer-term evolutionary dynamics of highly divergent, mosaic genomes and the extent to which genetic incompatibilities shape their adaptation. In yeast, pre-zygotic barriers are weak but post-zygotic barriers are strong due to mispairing of chromosomes during meiosis. By suppressing anti-recombination genes in meiosis, we generated a panel of 20 haploid recombinant hybrids from a cross between *Saccharomyces cerevisiae* and its sister species, *Saccharomyces paradoxus*. Across conditions, these hybrids are, on average, less fit than either parent and show broad phenotypic variation. Inheritance patterns of protein complexes in the hybrid genomes reveal no evidence of pairwise lethality but do support a model of pervasive weak negative genetic interactions in hybrid protein complexes. We show by laboratory evolution that each recombinant genome follows a distinct evolutionary trajectory, and a small subset of hybrid protein complexes and loci show hybrid-specific mutational targeting. Finally, we show that species-of-origin alleles can bias evolutionary outcomes by reshaping selection on interacting genes. Together, our results suggest that strong pairwise incompatibilities are rare, while weak, background-dependent incompatibilities are widespread and shape fitness and adaptation in hybrid genomes.

## INTRODUCTION

Understanding the genetic factors that establish and maintain species barriers is a central goal in evolutionary biology. Natural hybrids are widespread across taxa and offer powerful systems in which to assess the genetic basis of adaptation and speciation. Hybridization is a double-edged sword. On one hand, hybridization can disrupt interactions between coevolved genes, leading to incompatibilities that affect fitness or viability. This outcome of hybridization is well studied and is widely acknowledged as a key contributor to the maintenance and reinforcement of species barriers [1, 2]. On the other hand, hybridization can create or potentiate novel phenotypes absent from either parent [3, 4]. Although still debated [5], this idea raises the possibility that hybridization can facilitate rapid adaptation to novel environments [3]. However, how highly mosaic hybrid genomes respond to selection over many generations, and how incompatibilities shape their evolution, remains less well understood.

Interspecific hybridization is frequent between yeast in the genus *Saccharomyces* due to weak pre-zygotic barriers [6, 7]. Diploid F1 hybrids, formed from haploid strains of different species, are easily generated in the laboratory and are widespread in natural environments where they are valued for advantageous traits including altered metabolism, stress tolerance, and enhanced fermentation performance [8–11]. Numerous studies have shown that evolution of diploid hybrid genomes, both in nature and in the laboratory, is driven largely by loss of heterozygosity and genomic rearrangement [12–15].

While diploid F1 hybrids of *Saccharomyces* yeasts can be propagated mitotically and are often more fit than either parent due to hybrid vigor (heterosis), strong post-zygotic species barriers prevent the recovery of viable progeny following meiosis. Despite many attempts to identify Dobzhansky–Muller incompatibilities in yeast using genetic screens, chromosome-replacement experiments, genome-wide analyses of rare viable spores, and quantitative trait mapping in partially fertile crosses, no strong pairwise nuclear-nuclear incompatibilities have been identified [16–21]. While karyotype differences exist between some *Saccharomyces* species and these have been shown to contribute to species barriers [22, 23], the primary post-zygotic barrier is sequence divergence, which reduces meiotic viability by impairing proper chromosome pairing [24].

This post-zygotic barrier can be partially overcome by eliminating anti-recombination mechanisms that typically prevent mispairing in meiosis [24]. A recent study showed that placing two genes (*MSH2* and *SGS1*) under the control of a mitotic-specific promoter that is repressed in meiosis enables recovery of viable haploid recombinant hybrids (segregants) [25]. Unlike diploid F1 hybrids, these segregants fix allelic combinations to a single parental species at each locus, allowing the identification of interspecific nuclear-nuclear and nuclear-mitochondrial genetic incompatibilities [16, 21, 26, 27].

Here, we generate, phenotype, and evolve a panel of 20 segregants between *S. cerevisiae* and its sister species *S. paradoxu*s [28]. Across conditions, these segregants are, on average, less fit than either parent. By combining phenotyping with genetic mapping, we identify both Mendelian and quantitative traits in the segregants, including a defect in mother–daughter cell separation that maps to the mitotic regulator *AMN1*. To test for pairwise genetic incompatibilities, we classify the species-of-origin composition of all annotated protein complexes. Every two-subunit complex forms an interspecific (hybrid) assembly in at least one segregant, indicating that lethal pairwise incompatibilities are rare or absent; however, two-subunit hybrid complexes are underrepresented, indicating broad constraints on interspecific assembly. Finally, we evolve the segregants and their diploid derivatives for 10,000 generations and track their fitness gains and patterns of genomic change. In the evolved populations a small subset of genes and two-subunit complexes accumulate mutations more often than expected in hybrid genomes, identifying candidate targets of hybrid-specific selection; these include recurrent mutations in *HSP104* that arise in half of the hybrid lineages. We identify the *S. cerevisiae* allele of the prion factor *SUP35* as a potentiating variant that creates a strong selective pressure for mutations in its chaperone partner *HSP104*. Together, these results support a model in which pairwise negative genetic interactions are generally weak but pervasive, depressing fitness and shaping idiosyncratic adaptive trajectories in recombinant hybrid yeast genomes, and suggest that similar landscapes of weak, background-dependent incompatibilities may be common in other recently formed hybrid lineages.

## RESULTS

### Meiotic recombination produces balanced interspecific hybrid genomes

*S. cerevisiae* and *S. paradoxus* are sister species that diverged ∼5 million years ago. The two species are ∼85% identical at the amino acid level, yet the genomes are nearly completely syntenic (Supplementary Fig. 1). Of ∼6,600 annotated genes in *S. cerevisiae*, 5,366 have unambiguous one-to-one nuclear orthologs in *S. paradoxus*, and 99% of orthologs are not significantly different in size (Wilcoxon signed-rank, *p* < 0.001, Supplementary Data 1). These two species are separated by strong post-zygotic barriers due to improper chromosome pairing and segregation in meiosis, leading to spore viability of less than 1%.

We constructed prototrophic *S. cerevisiae* S288C and *S. paradoxus* CBS432 strains in which the mismatch repair gene *MSH2* and the helicase gene *SGS1* are under the control of the meiotically repressed *CLB2* promoter. Transient suppression of Msh2/Sgs1 during meiosis relaxes mismatch-repair–mediated anti-recombination between the highly diverged homologues, increasing crossovers and improving chromosome segregation. This manipulation increased spore viability to 23.5%, consistent with previous reports [25]. Using this approach, we isolated 20 segregants from five four-spore viable tetrads (Fig. 1a,b; Supplementary Fig. 2a). We confirmed by whole-genome sequencing that the segregants have mosaic nuclear genomes with nearly-equal contributions from both parents, and a uniparentally inherited mitochondrial genome (Fig. 1c; Supplementary Fig. 2b; Supplementary Data 1). Crossovers were markedly reduced. We find 17.6 crossovers per meiosis, ∼4.5 times fewer than the ∼70–90 crossovers that are typical in *S. cerevisiae* [29]. Approximately one quarter of chromosomes are uniparental throughout their length (Fig. 1d and Supplementary Fig. 2b), and the frequency in which a chromosome is observed without recombination is inversely proportional to chromosome size (Supplementary Fig. 2c).

**Fig. 1.**
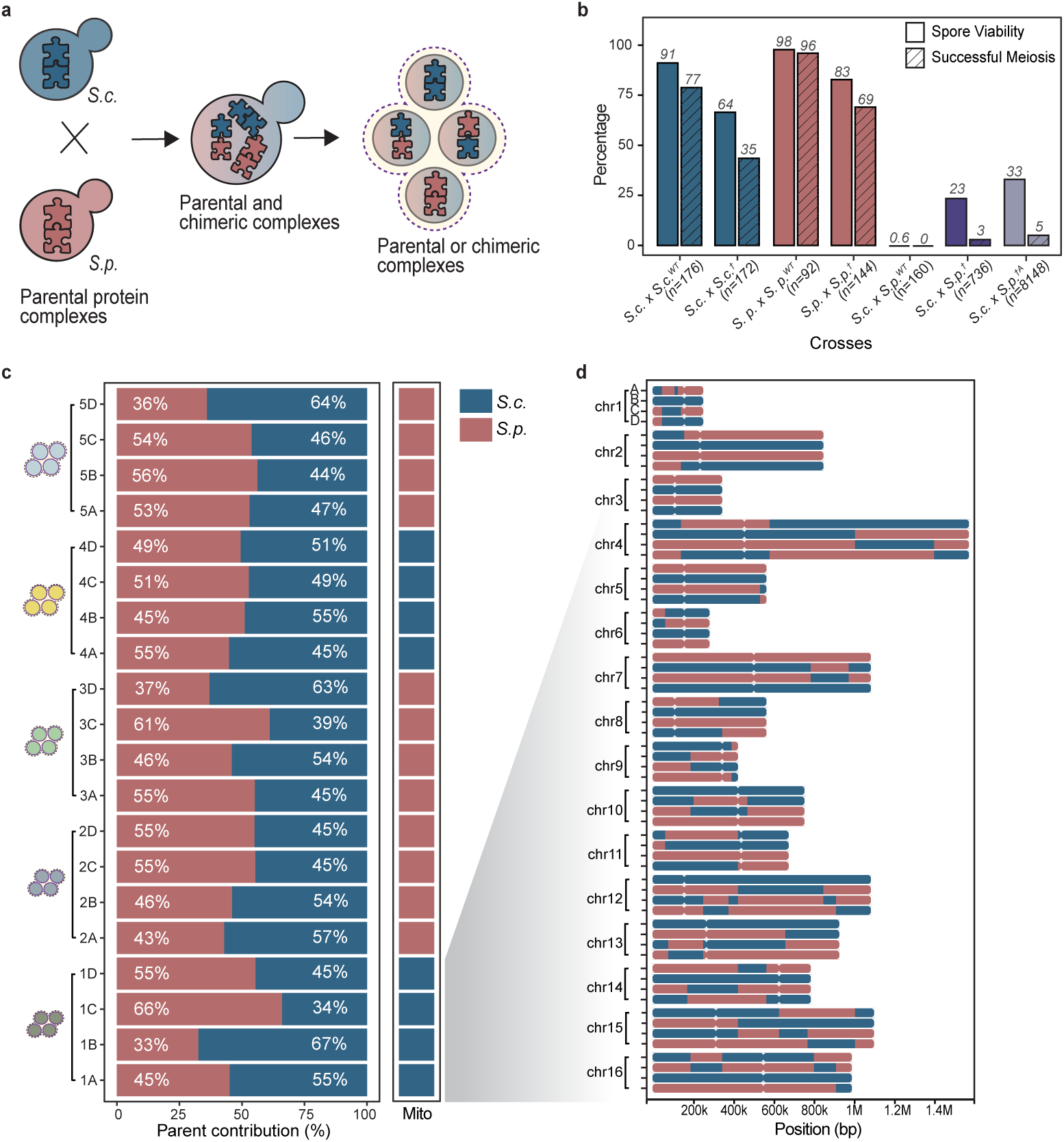
Construction and genomic architecture of recombinant haploid hybrids. **a,** Haploid recombinant hybrids (segregants) werer generated by placing the anti-recombination factors *SGS1* and *MSH2* under control of the meiosis-repressed *CLB2* promoter, transiently suppressing mismatch-repair–mediated anti-recombination during meiosis, increasing viable spore recovery and yielding genomes with mosaic nuclear ancestry. Recombination reshuffles parental alleles so that multi-subunit protein complexes can assemble in either parental or chimeric configurations; some chimeric assemblies are likely suboptimal and thus constitute potential targets of selection for compensatory change. **b,** Spore viability and successful meiosis for intra- and interspecific crosses of *S. cerevisiae (S.c.)* and *S. paradoxus (S.p.)*, with and without CLB2p-driven *SGS1/MSH2* suppression. Parentheses indicate total numbers of spores dissected. †, temporal suppression of *SGS1/MSH2*; A, data from *Bozdag et al., 2021*. **c,** Nuclear ancestry across 20 segregants (five tetrads), shown as the percentage contribution from *S. cerevisiae* (blue) and *S. paradoxus* (pink) in each genome. Adjacent bars indicate mitochondrial ancestry, which is uniparental and consistent within each tetrad. **d,** Chromosome-wide recombination maps for segregants A–D of Tetrad 1, showing balanced 2:2 segregation of parental alleles along each chromosome (colors as in panel c).

These recombinant genomes also give rise to both Mendelian and quantitative trait variation. Among these traits, clump formation is the most visually striking and reflects a parental difference in cellular morphology: *S. cerevisiae* primarily grows as dispersed single cells, whereas *S. paradoxus* forms small aggregates (Fig. 2a). Among 20 segregants, this phenotype segregated approximately 2:2 (Fig. 2a; Supplementary Fig. 3a). We identified a ∼222 kb region on Chromosome II whose segregation pattern matched the clumping phenotype in all but one spore. This interval contains 112 annotated genes, including *AMN1*, which has been previously implicated in mother–daughter separation [30]. Using CRISPR–Cas9, we exchanged the parental alleles of *AMN1* in both species and in four segregants derived from a single tetrad. The *S. paradoxus* allele (*AMN1*^368D^) inhibited mother–daughter separation leading to cell clumps, whereas the *S. cerevisiae* allele (*AMN1*^368V^) restored growth as single cells (Fig 2a). Thus, clumpiness segregates as a Mendelian trait in this interspecific cross, whereas most other phenotypes in the panel show quantitative, continuous variation.

**Fig. 2.**
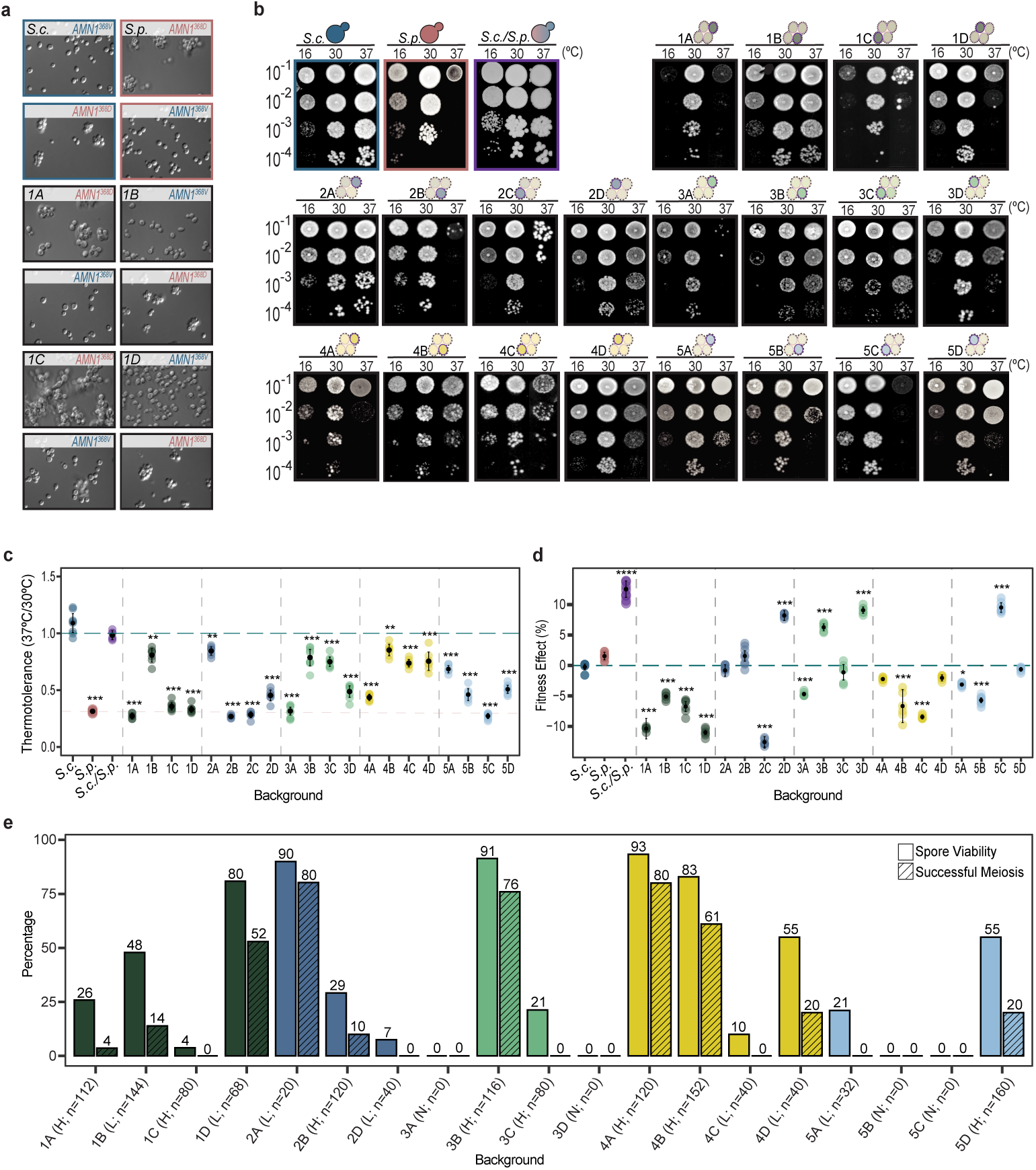
Segregants exhibit broad phenotypic diversity and reduced fitness. **a,** Clumping phenotype segregates 2:2 within each tetrad. Segregants carrying the *S. paradoxus (S.p.) AMN1*^368D^ allele display mother–daughter separa-tion defects, whereas those with the *S. cerevisiae (S.c.) AMN1*^368V^ allele separate normally. CRISPR–Cas9 allele swaps in parental strains and segregants confirm the causal effect of the *AMN1* allele. **b,** Spot-dilution assays at 30°C, 37 °C and 16 °C. Five-fold serial dilutions were plated and incubated for 2 days at 30 °C and 37 °C and for 3 days at 16 °C. Temperature-dependent growth profiles reveal both thermal sensitivity and emergent thermotolerance among segregants. The diploid F1 hybrid shows heterosis under all three conditions. **c,**Thermotolerance estimated from spot assays by quantifying spot intensities of 8 replicates per background across dilutions at 30 °C and 37 °C; growth at 37 °C was normalized to growth at 30 °C using the equal-average method. Asterisks mark thermotolerance values that differ significantly from the *S. cerevisiae* parent (Welch’s two-sample t-test on R = S_37_/S_30_; *p < 0.05*; \*\**p < 0.001*; data without asterisks are not significant). Error bars show mean ± s.e.m. (n = 8). **d,** Competitive fitness of 20 segregants relative to *S. cerevisiae*; *S. paradoxus* controls do not differ significantly from *S. cerevisiae.* Asterisks indicate significance (Z-test with Bonferroni correction: *p < 0.05*; \*\**p < 0.001*; data without asterisks are not significant). Error bars show mean ± s.e.m. (n = 4). **e,** Spore viability and frequencies of successful meiosis for segregant-derived diploids. Bars report the percentage of viable spores across all dissections (*n=* total spores dissected). Sporulation efficiency categories are marked H (high), L (low) or N (no spores). No diploid derivative was obtained for segregant 2C.

### Recombinant hybrids are on average less fit than either parent

We next asked how the segregants varied for quantitative traits affecting growth and reproduction. We quantified traits linked to fitness in the segregant panel, including thermotolerance, growth rate, competitive fitness, and sporulation efficiency (Supplementary Data 2). Consistent with previous observations [31], *S. cerevisiae* exhibits greater thermotolerance than *S. paradoxus*. The diploid F1 hybrid exceeded the growth of both parents. By contrast, the 20 segregants showed a broad range of temperature responses, with most segregants growing well at 30 °C, but failing to form robust colonies at 37 °C (Fig. 2b). Quantification of growth at 37 °C relative to 30 °C shows that only a few segregants approach the thermotolerance of *S. cerevisiae*. Segregants 2A, 4B, and 1B reached ∼80–85% of their 30°C growth at 37 °C, but remained significantly below *S. cerevisiae* (∼75–80% of the parental value; two-sample t-tests vs *S. cerevisiae*, *p* < 0.0001 for each) (Fig. 2c). Similar variability was also observed at 16 °C. Direct growth-rate measurements corroborated these findings, showing intermediate phenotypes with a marked decline at 37 °C, consistent with the spot assay results (Supplementary Fig. 3b, c). Together, these results show a broad range of temperature responses, with the strongest divergence at high temperature, identifying thermotolerance as an important varying phenotype in the segregants.

To quantify overall performance, we next measured competitive fitness at 30 °C in rich glucose medium. At this temperature, the two parental species show comparable growth, whereas the diploid F1 hybrid displays clear heterosis. In contrast, most segregants are less fit than one or both parents, and only a few exceed both (Fig. 2d). Competitive fitness correlates moderately with growth rate (r = 0.56), but with several strong outliers (Supplementary Fig. 3d), consistent with contributions from additional traits beyond simple growth rate.

Given the pronounced variation in vegetative growth, we next asked whether sexual fitness is similarly affected, and measured sporulation efficiency for each segregant. We constructed a segregant-derived diploid (a homozygous diploid formed by mating the segregant to itself). Unlike vegetative growth, sporulation requires accurate homolog pairing, recombination, and chromosome segregation; defects therefore implicate failures in core meiotic processes. We observed wide variation in both sporulation efficiency and spore viability (Fig. 2e), consistent with reduced meiotic fitness in hybrid genomes. Both growth and sporulation are sensitive to mitochondrial function, and nuclear–mitochondrial incompatibilities have been described previously in Saccharomyces yeasts [20]. However, despite several segregants carrying the previously reported incompatible combination of *S. paradoxus* mtDNA and the *S. cerevisiae MRS1* allele [19], none showed respiratory defects (Supplementary Fig. 3e).

### Weak selection acts against hybrid protein complexes

Having established that many recombinant genotypes show reduced performance across multiple environments, we next asked whether any signal of incompatibility could be detected at the level of protein–protein interactions. One plausible source of such incompatibilities is the misassembly of protein complexes when subunits from different species are forced to interact. If chimeric complexes were strongly deleterious, we would expect surviving recombinant genomes to be depleted of hybrid assemblies and enriched for uniparental complexes, especially among small complexes where each subunit contributes a large fraction of function.

We curated a set of 628 conserved protein complexes (Supplementary Data 3) and, for each of the 20 segregants, classified every complex as “hybrid” (containing ≥1 subunit from each species) or “uniparental” (all subunits from a single species). As expected, smaller complexes (2–4 subunits) were hybrid in ∼62% of complex–segregant combinations, whereas larger assemblies (≥5 subunits) were hybrid in ∼96%. (Supplementary Fig. 4a). Across all 13,040 complex–segregant combinations, the probability that a complex assembles as a hybrid increases by ∼2.3-fold with each additional subunit (logistic regression, *p < 10^-16^*), consistent with a combinatorial expectation that larger assemblies are almost always chimeric. Because each subunit contributes a larger fraction of function in small complexes, any constraint on interspecies interactions should be most evident in two-subunit assemblies.

To test whether hybrid protein complexes are under- or overrepresented relative to a neutral expectation, we compared the observed frequencies of hybrid and uniparental two-subunit protein complexes against a null model that takes into account that spores from a single tetrad are not independent. We focused on two-subunit complexes where hybrid and uniparental configurations have equal baseline probability, excluding protein complexes where both subunits are encoded by genes on the same chromosome. Notably, every two-subunit hybrid complex was recovered in at least two independent segregants, showing that there are no pairwise lethal incompatibilities. We observe, however, a slight skew toward uniparental complexes in the overall distribution, consistent with weak selection against hybrid assembly (Fig. 3a). We performed 10,000 randomizations of chromosomes within each of the five tetrads and recalculated the distribution of hybrid complexes. Fewer than 1% of the reshuffling trials produced a greater excess protein complexes observed as hybrids in only 4 or 6 of the segregant genomes. In addition, protein complexes that occur by chance in the majority of segregant genomes are more likely to be nonessential (Fig. 3b).

**Fig. 3.**
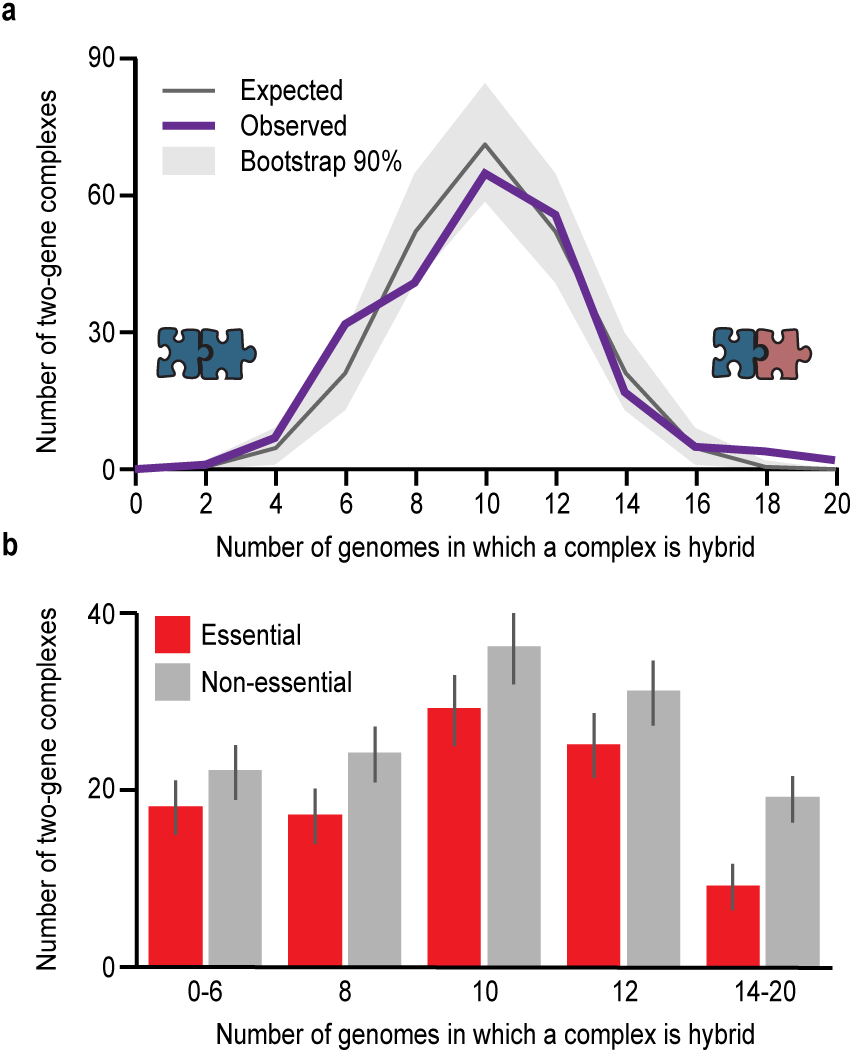
Two-subunit protein complexes show a slight but significant skew towards uniparental origin. **a,** For each of 230 two-subunit complexes, we calculated the distribution of the number of segregant genomes in which the complex assembles as hybrid versus uniparental. We excluded complexes that are genetically linked on the same chromosome and one complex where the parent of origin is ambiguous. The observed distribution deviated significantly from the expected distribution (*p < 0.001*, Chi-Sq). To assign significance, we performed 10,000 bootstrapping iterations by computationally shuffling the cromosomes within each of the five tetrads and recalculating the distribution of hybrid complexes. The bootstrapped data matched the expected distribution: 91.2% of resampling experiments had chi-squared *p-values > 0.01*, and only 8 of the 10,000 resampling experiments showed a stronger deviation from the expected distribution than the observed data. **b,** Essential two-subunit complexes are observed less frequently as hybrid complexes relative to non-essential complexes. We define essential complexes as ones where either protein is classified as essential by the *Saccharomyces* Genome Database (yeastgenome.org). Error bars are the standard deviation expected due to binomial sampling.

### Hybrid genomes follow unique evolutionary trajectories

Having established that many segregants are compromised, we next asked how selection acts on loci affected by weak incompatibilities. We performed a 10,000-generation long-term evolution experiment with 8 replicate populations for each of the 20 segregants, the two parental strains, and their segregant-derived diploid (or polyploid). We measured fitness by competition against a fluorescently labeled haploid or diploid version of *S. cerevisiae*, respectively. Initial fitness of segregants and their segregant-derived polyploid was only moderately correlated (r = 0.51; Fig. 4a), with several backgrounds either gaining or losing fitness following ploidization. Segregant-founded lines evolved heterogeneously: some improved steadily, others diverged among replicates, demonstrating strong genome background effects on adaptive trajectories. In general, we find that low-fitness founders had the most rapid initial rate of adaptation (Fig. 4b,c).

**Fig. 4.**
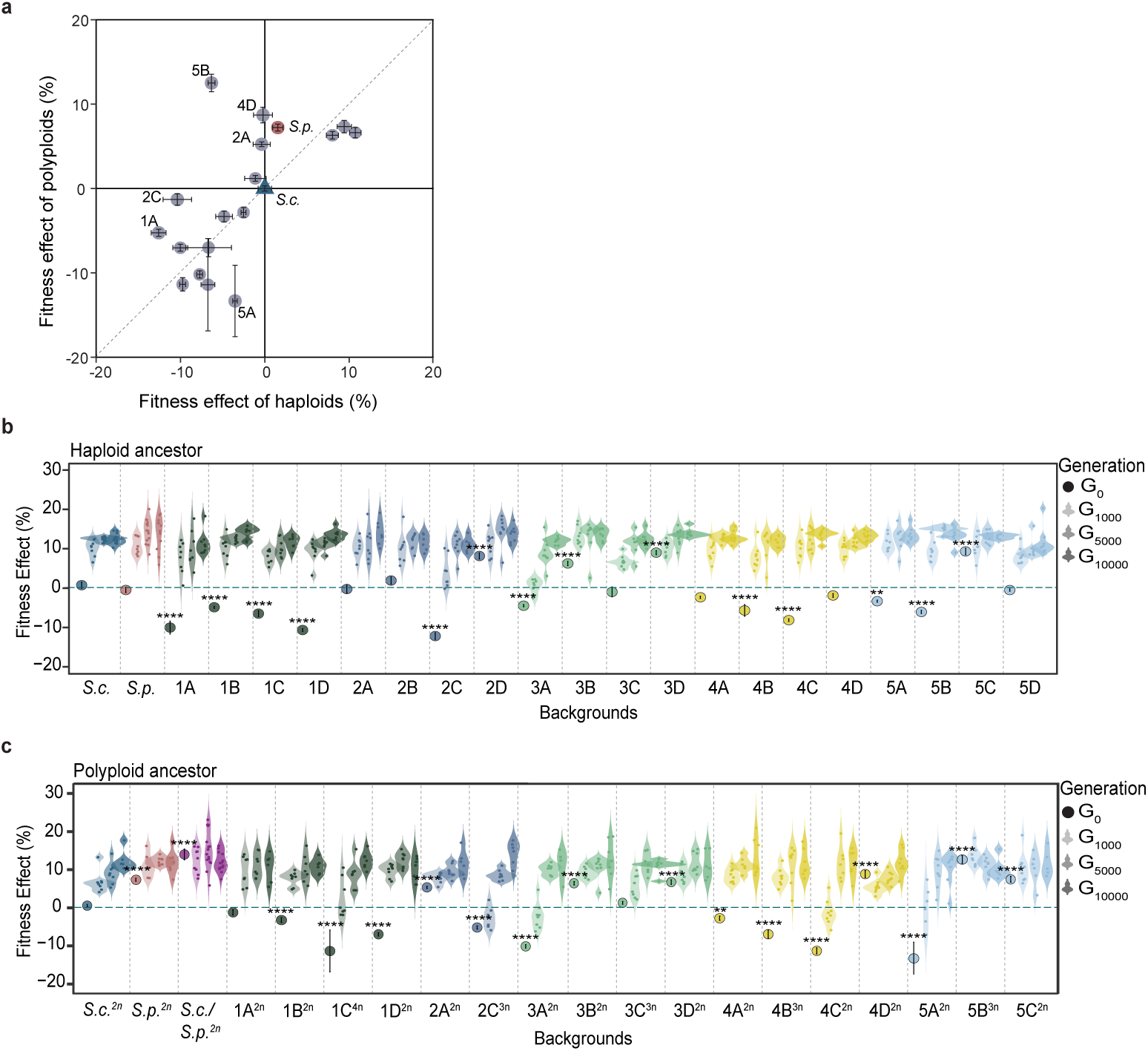
Long-term fitness trajectories of segregants and their derivatives. **a**, Relationship between selection coefficients of segregants as haploids (x-axis) and as segregant-derived diploids (y-axis) at 30 °C. Pearson’s r = 0.51. Points corresponding to segregants 1A, 2A, 2C, 4D, 5A and 5B differ by more than 0.05 in their haploid versus diploid selection coefficients. **b,** Relative fitness of haploid parental strains and segregants at generations 0, 1,000, 5,000 and 10,000. Fitness is expressed relative to the *S. cerevisiae* ancestor. Circles mark ancestral fitness (generation 0); violin widths reflect the distribution of evolved replicate populations per background. Asterisks denote significance (Z-tests with Bonferroni correction; \**p < 0.05*; \*\*\**p < 0.001*; data without asterisks are not significant). **c,** As in b, but for segregant-derived diploid and polyploid lines. Ploidy states are annotated as 2n (diploid), 3n (triploid) and 4n (tetraploid).

Although evolution occurred at 30 °C, we used thermotolerance at 37 °C as a proxy for proteostasis and stress-buffering capacity, because negative genetic interactions are often exacerbated at elevated temperature. If evolution in hybrids resolves underlying conflicts in protein homeostasis, we would expect correlated gains in growth at 37 °C even without direct selection at that temperature. Among the parental lines, *S. cerevisiae* already grew robustly at 37 °C and showed little change over evolution (gen 5,000 *p = 0.70*; gen 9,000 *p = 0.19*), whereas *S. paradoxus* remained poorly thermotolerant and even showed a transient reduction at generation 5,000 (*p = 0.002*), with one replicate improving (Supplementary Fig. 5). Segregant-founded populations, in contrast, displayed heterogeneous trajectories. Comparing evolved populations to their own ancestors, 10/20 hybrids showed a significant decrease in thermotolerance at generation 5,000, and across all lineages the change in thermotolerance between generation 0 and 5,000 was on average −0.13 ± 0.23 (mean ± s.d., *p < 10^-12^*), indicating a substantial global reduction early in evolution. By generation 9,000, 7/20 hybrids remained significantly less thermotolerant than their ancestor, but globally the mean change had attenuated to −0.05 ± 0.59 and was no longer significantly different from zero (*p = 0.26*). At the lineage level, “strong improvers” were relatively rare (∼20% of populations) and were strongly enriched in a small subset of backgrounds (notably 5C and 5D). Thus, despite frequent modest losses, long-term evolution at 30 °C did produce clear increases in thermotolerance in a subset of hybrids, showing that high-temperature performance can improve even without direct selection.

### Genome background affects the evolutionary trajectories of recombinant hybrids

To identify the genetic targets of selection, we sequenced evolved populations from segregant-derived diploid/polyploid lines at generation 2,400 and from segregant-founded haploid lines at generation 5,000. In total, we analyzed 336 independently-evolved populations, including 275 hybrid populations (Supplementary Fig. 7a; Supplementary Data 4). Across the 163 diploid/polyploid-founded populations, we identified 14,246 variants, most of which (87.3%) were called as heterozygous (Supplementary Fig. 6a). Across the 173 haploid-founded populations, we identified 21,933 variants (Supplementary Fig. 6b). Autodiploidization is a common outcome of laboratory evolution experiments [32, 33], and our haploid-founded lines were no exception: 78.5% of mutations in these populations are heterozygous (Supplementary Fig. 6a,c). We confirmed by flow cytometry that >90% of our haploid-founded lines underwent autodiploidization. Interestingly, 9 out of the 14 populations founded from segregants 4C (4/8) and 5D (5/6) remained haploid throughout 5,000 generations (Fisher’s exact *p = 0.003* and *p = 0.0002*, respectively; Supplementary Fig. 6d).

Across hybrid genomes, we detected 29,650 mutations (38.6% intergenic, 32.2% missense, 15.0% synonymous, 11.4% frameshift, 2.6% nonsense), comparable to parental distributions (Supplementary Fig. 7b). Because we replaced the native promoter of *MSH2*, we anticipated a slight elevation in mutation rate. Through 5,000 generations, we observe an average of 110 ± 66 mutations per lineage (Supplementary Fig. 7c,d), above the ∼30 mutations we expected based on prior evolution experiments [32, 34, 35]. This excess is driven largely by frameshift mutations, which are ∼4-fold more frequent than in earlier studies. Interestingly, populations founded from two hybrid backgrounds (3C and 5B) have an excess of mutations relative to the other segregants, indicating a potential mutator phenotype (Supplementary Fig. 7c).

We identified loci accumulating more nonsynonymous mutations than expected by chance. Among 13,283 nonsynonymous mutations (Fig. 5a), of which 9,139 are nuclear missense mutations (Supplementary Fig. 8a), shared targets—*IRA2*, *UPC2*, and *PTR2*—were enriched for nonsynonymous mutations in both hybrids and parents (Supplementary Fig. 8b,c), whereas *HSP104* and *BSC1* are strongly enriched only in hybrids despite a small number of mutations in the parental backgrounds (Supplementary Fig. 8d,e).

**Fig. 5.**
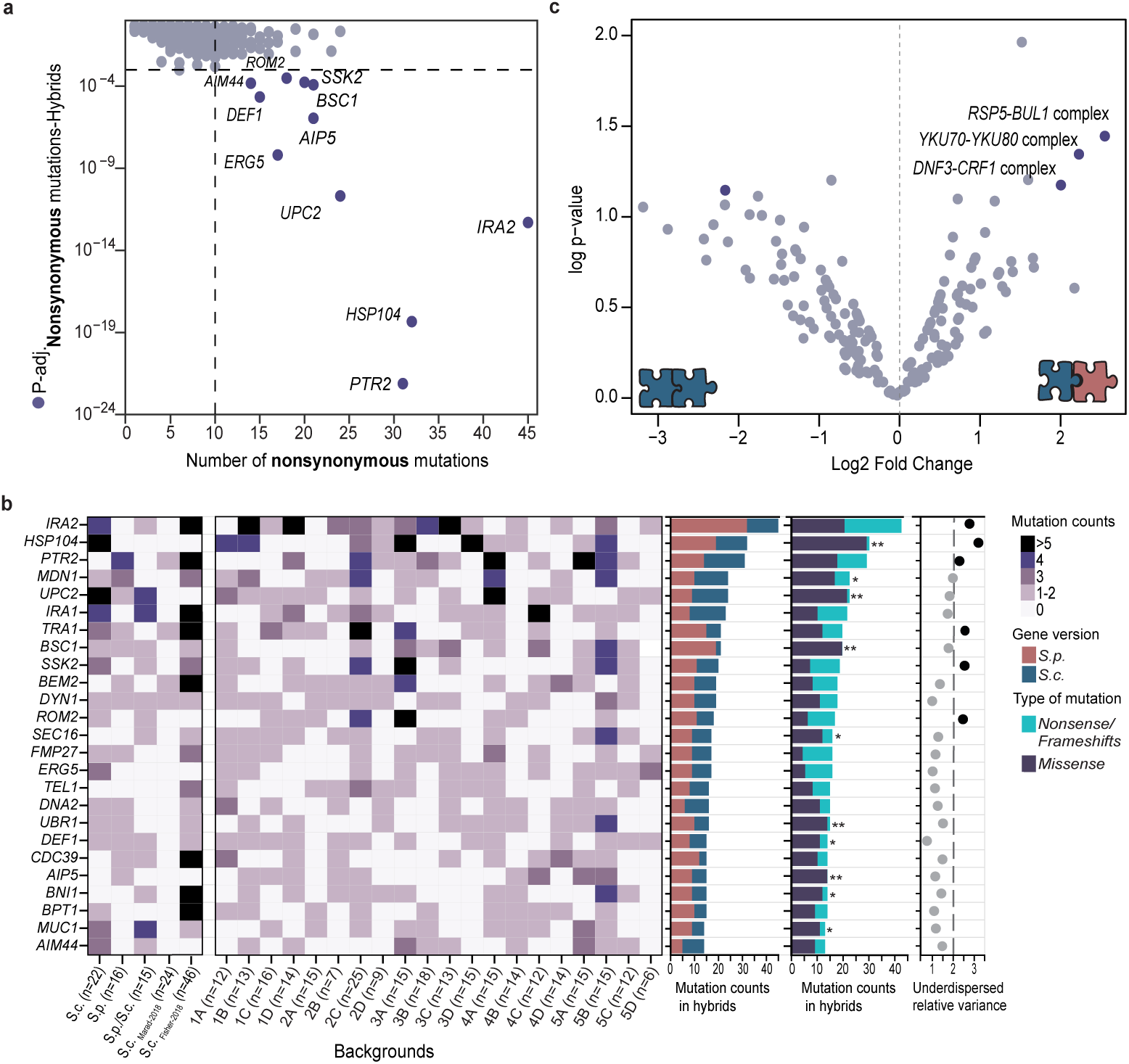
Gene- and complex-level targets of selection in evolved hybrid genomes. **a,** Gene-level enrichment of nonsynonymous coding mutations across ∼5,800 CDSs in hybrids, evaluated under a Poisson expectation. The x-axis shows the number of nonsynonymous mutations per gene; the y-axis shows the probability of observing that number by chance, weighted by the average CDS length across species. Genes exceeding the multiple-testing threshold (Benjami-ni–Hochberg–adjusted *p < 10^-4^*) and with at least 10 nonsynonymous mutations are highlighted in purple. **b,** Recurrently mutated genes. Left, heat map of the top 25 genes ranked by total nonsynonymous counts across hybrid populations, parental controls and published datasets (Fisher et al., 2018; Marad et al., 2018) (darker shading indicates higher mutation counts). Centre, bar plots partitioning counts by allele of origin—*S. paradoxus (*pink, *S.p*.) versus *S. cerevisiae* (blue, *S.c*.). Right, mutation class and selection signal: bars show missense (purple) versus nonsense/frameshift (blue) mutations, with overlaid posterior probabilities that selection favors non-loss-of-function; asterisks denote significance (\**p < 0.001*; *p < 0.05*). Prior probabilities were based on seven independent yeast evolution experiments (Martínez & Lang, 2023). Adjacent dot plot: variance (underdispersion) scores quantify deviation from neutral expectations; genes with scores ≥ 2 (dashed line) show repeatable, allele-specific targeting consistent with selection. **c,** Enrichment of mutations in 251 two-subunit protein complexes in hybrids relative to parental controls. The x-axis is the log2 fold-change in mutation enrichment and the y-axis is –log10 *p*. Complexes that pass both the effect-size threshold |log2 fold-change| ≥ 2 and the significance threshold –log10 *p ≥ 1.1* are highlighted in dark purple.

Each hybrid genome is a unique mosaic of genes from *S. cerevisiae* and *S. paradoxus*. In the absence of interspecies genetic incompatibilities, selection is expected act similarly on each hybrid genome, leading to parallel evolutionary outcomes. However, if genomic mosaicism leads to widespread and background-specific genetic incompatibilities, evolutionary outcomes will be idiosyncratic, with selection acting on different loci in each hybrid background, even when the hybrids experience identical environments. To distinguish between these scenarios, we measured the coefficient of variation for the number of mutations observed in a given gene across the parental and hybrid backgrounds. Among the 25 most frequently mutated genes, 6 are significantly under-dispersed, where mutations are clustered more than expected by chance in specific backgrounds (Fig. 5b).

Most genes that were identified as targets of selection carry mutations in both parental alleles, with some locus-specific skews (*IRA1*, 33/46 mutations in the *S. paradoxus* allele; *BSC1*, 19/21 mutations in the *S. paradoxus* allele). To infer whether selection favors complete loss-of-function or alteration/attenuation-of-function, we applied a likelihood-based approach to the observed mutational spectrum at each target gene (Fig. 5b). *IRA2* and *PTR2* accumulate missense, nonsense, and frameshift mutations, consistent with selection for loss-of-function. In contrast, *HSP104*, *UPC2*, and *BSC1* accumulate mostly missense mutations, suggesting that selection is acting to alter or attenuate the function of these genes.

Finally, we asked whether hybrid protein complexes are preferential targets of compensatory evolution. Among two-subunit complexes, deviations from the null expectation were rare: only three complexes showed significant hybrid-enriched targeting: *RSP5–BUL1*, *YKU70–YKU80*, and *DNF3–CRF1* complex (Fig. 5c). Together, these results indicate that hybrid-specific selection on complex assembly is infrequent, supporting a model in which interspecies protein interactions are broadly tolerated, with only subtle, complex-specific constraints.

### The *S. cerevisiae* allele of *SUP35* imposes selection on *HSP104*

*HSP104* was among the most frequently mutated genes in the hybrid populations and is strongly underdispersed (32 nonsynonymous mutations; underdispersion score = 3.25; *p < 10^-13^*) across backgrounds (Fig. 5B). Mutations in *HSP104* were observed in *S. cerevisiae* but not in *S. paradoxus* populations despite 99.3% amino acid identity, suggesting background-dependent selection. A genome-wide scan of parental ancestry revealed a single overrepresented region: nearly all hybrids with *HSP104* mutations retained a conserved *S. cerevisiae* haplotype (∼318 kb) on Chromosome IV containing the interactors *SUP35* (96.5% identity) and *HSP42* (85.7% identity). The only exception (5A) lacked this block but carried a mutation in *HSP42* (Fig. 6a). Additional *SUP35* mutations (4B, 4D) further suggest that this gene is likely imposing selection on *HSP104*.

**Fig. 6.**
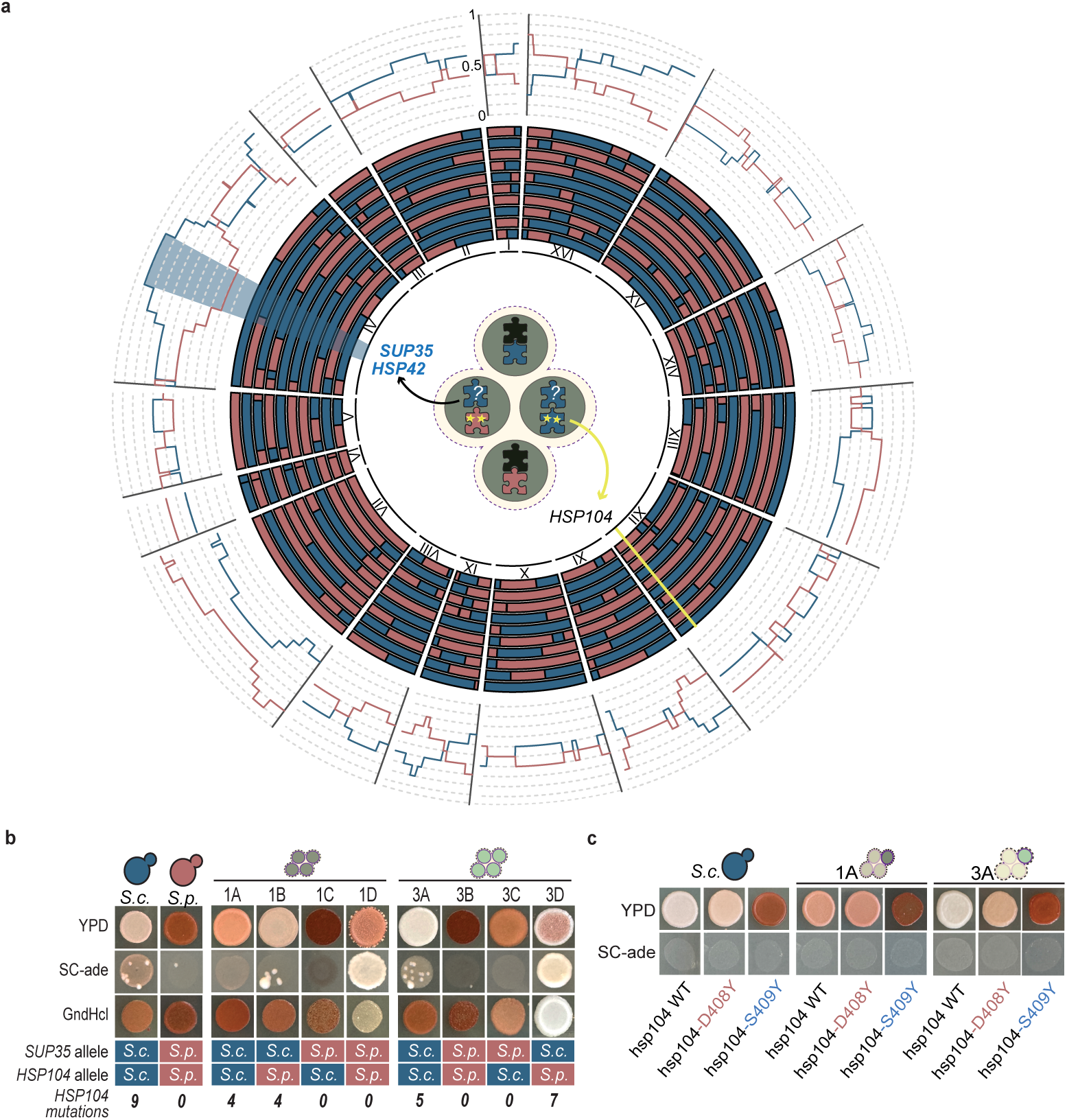
An *S. cerevisiae* chr IV block containing *SUP35* co-segregates with recurrent *HSP104* mutations and predicts their occurrence in hybrid segregants. **a,** Circos maps of segregants that gave rise to *HSP104* mutations in evolved populations (∼50% of segregants). Hybrids whose populations acquired *HSP104* mutations consistently retain an *S. cerevisiae* (blue, *S.c*.) segment on chromosome IV; the only exception (5A) carries the corresponding *S. paradoxus* (pink, *S.p*.) block. This interval spans *HSP42* and *SUP35*, known physical interactors of Hsp104. **b,** Prion phenotyping of parents and segregants on YPD, SC–Ade and 5 mM guanidine-HCl, using the ade1-W244* translational readthrough reporter (non-prion colonies red; prion colonies pink/white). The *S. cerevisiae* parent exhibits a weak [PSI+] state, whereas the *S. paradoxus* parent is [psi-]. Segregants from tetrads 1 and 3 span parental-like and intermediate prion states, indicating differences in prion strength across hybrid backgrounds. The table below summarizes allele origin and mutational load: each square indicates the allele of origin for *SUP35* and *HSP104*, and the numbers indicate how many *HSP104* mutations arose in evolved populations founded from each segregant. Prion phenotyping was performed with four biological replicates per genotype. **c,** Functional reconstructions of evolved Hsp104 variants. Two recurrent mutations—Hsp104-D408Y (c.1222G>T; p.Asp408Tyr), which arose in a population founded from segregant 1B carrying the *S.p*. *HSP104* allele (pink), and Hsp104-S409Y (c.1226C>A; p.Ser409Tyr), which arose in a 3A-founded population carrying the *S.c*. HSP104 allele (blue)—were engineered into the *S.c. HSP104* allele in the *S. cerevisiae* parent and in segregants 1A and 3A. Prion phenotypes of reconstruction strains were assayed as in b; two independent transformants were tested for each genotype.

*SUP35* encodes a prion-forming translation termination factor. We assayed the prion state using an *ade1*-W244* translational readthrough reporter, in which non-prion colonies are red and prion colonies are pink/white. The *S. cerevisiae* parent exhibits a weak prion phenotype, whereas *S. paradoxus* is non-prion, and the recombinant hybrids span the full range from strong prion (1B, 3A) to weak prion (1A ,1D, 3C, 3D) to non-prion (1C, 3B) (Fig. 6b). Inhibiting Hsp104 activity with GdnHCl shifted most lines toward the non-prion state, but two hybrids (3D, 1D) remained white and grew robustly on –Ade, indicating genetic restoration of adenine biosynthesis rather than simple prion curing. To test how evolved *HSP104* mutations affect prion state, we reconstructed two evolved mutations in the *S. cerevisiae* parent and in two segregants (1A and 3A) that carry both the *S. cerevisiae* SUP35 allele and the *S. cerevisiae HSP104* allele; direct editing of the *S. paradoxus HSP104* allele was unsuccessful. The *hsp104-*D408Y substitution, which originally arose in a population founded from segregant 1B where the resident allele is *S. paradoxus*, caused only modest shifts in prion strength when introduced into *S. cerevisiae* and the segregants. By contrast, *hsp104*-S409Y, which arose in a 3A-founded population carrying the *S. cerevisiae* allele, consistently cured the prion phenotype when reconstructed in the *S. cerevisiae* parent and in both segregants (Fig. 6c). Overall, these results show that the *S. cerevisiae SUP35* haplotype redirects selection onto *HSP104* and that the consequences of *HSP104* mutations for prion state are strongly conditioned by hybrid genetic background.

## DISCUSSION

Meiosis fails in interspecific yeast diploids due to defects in chromosome pairing and segregation [24]. Eliminating or suppressing anti-recombination genes partially overcomes this strong post-zygotic barrier [6]. Using this strategy, we isolated 20 viable recombinant hybrids between *S. cerevisiae* and *S. paradoxus*, whose nuclear genomes are a balanced mosaic of the two evolutionarily divergent yet syntenic parental genomes. These segregants provide a powerful system to dissect how hybridization reshapes the fitness landscape and redirects the genetic targets of selection. The segregants display a range of phenotypic variation, from clear 2:2 segregation of a cell separation phenotype that maps to the *AMN1* locus, to quantitative, continuous variation in growth rate, thermotolerance, and sporulation efficiency. While some segregants have higher fitness, on average, they are less fit than either parent across tested conditions, consistent with fitness-reducing genetic interactions arising when coadapted genomes are recombined.

All hybrid two-protein complexes are observed at least once, establishing that all two-protein hybrid complexes are compatible with viability. This is in line with proteomics studies in diploid interspecific hybrids, which show that chimeric protein complexes form with similar stoichiometries to uniparental complexes [11, 36]. Yet we also find that two-protein complexes show a slight but significant overall bias toward uniparental composition in the viable hybrids. Together, these results suggest weak but pervasive selection to preserve intra-species coadaptation, rather than strong selection against a small number of interspecific assemblies. This is consistent with previous work that also found no evidence for classic Bateson–Dobzhansky–Muller incompatibilities between *Saccharomyces* yeasts [20, 37, 38]. Unlike nuclear-nuclear incompatibilities, several nuclear-mitochondrial incompatibilities are known to exist [16, 21, 39]. Surprisingly, however, we do not observe the previously identified incompatibility between an *S. cerevisiae* nuclear-encoded gene, *MRS1*, and the *S. paradoxus* mitochondrial genome [16], likely reflecting strain-specific differences between the *S. paradoxus* background used.

These initial incompatibilities shape subsequent adaptive trajectories over the course of 10,000 generations of laboratory evolution. Because each segregant lineage began with a unique mosaic of *S. cerevisiae* and *S. paradoxus* alleles, with a different set of weak negative interactions, each background adapted on a different fitness landscape. Most genetic targets of selection are not determined simply by the species-of-origin of the gene itself. The one exception is *BSC1,* which was observed as a significant target of selection only in hybrids that carried the *S. paradoxus* version of this gene (Fig. 5b). *BSC1* is a highly diverged, repeat-rich locus. The *S. paradoxus* version of the gene is significantly larger than the *S. cerevisiae* version due to a fusion event (Supplementary Fig. 1b, Supplementary Fig. 9b,c). This gene is also highly variable within *S. cerevisiae* [40]. The mutational signatures are consistent with intra-genic rearrangements between repeat sequences, rather than *de novo* SNPs (Supplementary Fig. 10b,c), suggesting that *BSC1* is a structurally labile locus whose involvement in adaptation depends on hybrid background.

To identify genes for which selection acts differently across segregants, we calculated the coefficient of variation for the number of times a mutation arose in each of the 20 hybrid founders. *HSP104* had the highest dispersion: the 32 *HSP104* mutations arose in just 10 of the 20 hybrid backgrounds, and in the *S. cerevisiae* but not the *S. paradoxus* parent. Interestingly, each of the five tetrads from which the hybrid lineages were derived show 2:2 segregation of the propensity to acquire *HSP104* mutations, suggesting that a single locus from the *S. cerevisiae* is imposing selective pressure on *HSP104*. We identify the *S. cerevisiae* version of *SUP35* as the likely potentiating gene and show that the *S. cerevisiae* version of Sup35 has a greater potential to form prion aggregates, which mutations in *HSP104* resolve.

Thermotolerance evolution in these segregants provides an additional window into how such incompatibilities are remodeled. We initially treated growth at 37 °C as a proxy for the burden of temperature-sensitive genetic interactions, yet long-term evolution at 30 °C produced highly heterogeneous thermotolerance trajectories across segregant backgrounds. Most hybrids showed modest losses or no net change in thermotolerance, whereas a minority of backgrounds produced “strong improver” lineages whose final high-temperature growth rose well above their ancestral distribution. This pattern suggests that adaptation often proceeds without fully resolving temperature-sensitive incompatibilities and that only specific allelic combinations permit compensatory evolution that improves both baseline fitness and proteostasis-related stress tolerance.

Our results support the model of weak but pervasive genetic incompatibilities in hybrid protein complexes. This model predicts that selection should act more strongly on chimeric protein assemblies—complexes composed of subunits derived from both *S. cerevisiae* and *S. paradoxus*—than on uniparental complexes. Consistent with this, we identify three two-subunit complexes—*RSP5*–*BUL1* (ubiquitin ligase), *YKU70*–*YKU80* (telomere end-binding), and *DNF3*–*CRF1* (P4-ATPase)—that show excess mutational targeting when assembled in hybrid form, indicating potential selective pressures acting on these chimeric protein complexes in hybrid genomes. Notably*, YKU70–YKU80* has previously been shown to evolve rapidly and to generate incompatibilities even between *S. paradoxus* populations [41, 42], reinforcing the view that divergence in these complexes can promote hybrid dysfunction.

Overall, our findings demonstrate how hybridization can both constrain and diversify adaptation; weak, distributed incompatibilities depress baseline fitness and subtly bias which genes and complexes are repeatedly targeted, while genome mosaicism and background-specific interactions generate diverse, lineage-specific evolutionary trajectories.

## METHODS

### Strain construction

All strains derive from prototrophic and heterothallic versions of *S. cerevisiae* S288C/DBY12000 and *S. paradoxus* CBS432. We replaced the native promoters of *SGS1* and *MSH2* using the p*CLB2*::3xHA::KanMX cassettes from Ref. [25] to create our parental strains yGIL2099 (*MAT***a**, *S. cerevisiae*) and yGIL2279 (*MAT*α, *S. paradoxus*). The diploid F1 hybrid yGIL2283 was generated by crossing these parents. Viable spores from yGIL2283 were isolated by tetrad dissection and archived. Strains were sporulated in SPO++ (1.5% potassium acetate with supplements) for ∼6 days at room temperature, ascus walls digested with Zymolyase, and tetrads dissected. We performed 184 dissections of yGIL2283 to isolate five four-spore tetrads (20 segregants). Segregant-derived diploids were obtained by transient expression of a galactose-inducible *HO* in each haploid segregant. We verified mating type by PCR and crossed complementary mating types to form homozygous diploids. Ploidy was assessed by testing for sporulation and by quantifying DNA content by flow cytometry. For three segregants (1C, 2C and 5B), standard HO induction was inefficient, so we extended induction overnight in galactose. For 1C and 5B, we recovered both diploid and polyploid derivatives (tetraploid and triploid, respectively), whereas for 2C only a triploid derivative was obtained. All of these segregant-derived lines were included in the long-term evolution experiment.

### Growth/thermotolerance phenotyping (spot assays)

Overnight cultures were arrayed in 96-well plates and serially diluted 1:32 in four steps (undiluted, 1:32, 1:1,024, 1:32,768; ∼10^-5^ overall). Dilutions were prepared on a Biomek Independent Span-8, and 5 µL from each dilution was pinned to YPD agar using the Biomek multichannel head. Plates were incubated 2 days at 30 °C or 37 °C, and 3 days at 16 °C, scanned under fixed settings, and spot intensities quantified in Fiji/ImageJ with local background subtraction. For each sample, intensities across dilutions were averaged to produce S_30_ and S_37_; thermotolerance was defined as R=S_37_/S_30_ , which integrates growth across dilutions with equal weight. The same assays were performed for evolve populations generations 0, 5,000, and 9,000; trajectories connect per-population R across timepoints, with generation-0 replicates averaged per segregant founder. Within each hybrid background, we then performed paired, two-sided *t*-tests on ΔR(R_5000_ – R0, R_9000_-R_0_) across lineages to test whether evolved thermotolerance differed from the ancestral state. We additionally classified “strong improver” lineages as those whose final thermotolerance at generation 9,000 satisfied R_equal,9000_ > µ_0_ + 3a_0_, where µ_0_ and a_0_ are the mean and standard deviation of ancestral R across lineages for that segregant founder.

### Growth curve analysis

Yeast growth in YPD was monitored at 17 °C, 30 °C, and 37 °C using a Tecan Infinite 200 Pro plate reader in Corning 96-well flat-bottom transparent polystyrene plates (353072). OD600 was recorded at fixed intervals with orbital shaking (10 s, 1 mm amplitude) before each read and multiple reads per well (2×2 grid). Media-only wells were used for background subtraction. Background-subtracted curves were fit to a Gompertz growth model using scipy.optimize.curve_fit to estimate key parameters, including maximum specific growth rate (µ_max), carrying capacity (A), and lag time (λ).

### Long-term experimental evolution

We founded 384 lineages (4 × 96-well plates) from saturated YPD cultures with genotypes randomized by position (Supplementary Data 3). Each day at the same time, cultures were serially diluted 1:32 twice into fresh YPD supplemented with ampicillin and tetracycline, following the same dilution and passaging regimen described in Martínez et al. (2022), yielding ∼10 generations per day and propagating populations for up to 10,000 generations.

### Competitive fitness assays

Fitness was measured by head-to-head competition against a fluorescent reference (yGIL519 (*MAT***a**), yGIL699 (*MAT*α), and yGIL702 (*MAT***a**/α) as described previously [43]. For flow cytometry (BD FACSCanto II), 4 µL of culture was diluted into 60 µL PBS. A minimum of 30,000 events was recorded per sample. The selection coefficient (per 10 generation) was estimated as the slope of ln(p_ref/p_population) versus generations. For segregants at generation 0, four biological replicates were assayed; evolved populations were assayed once per lineage. Statistical analyses were performed in R. Differences in mean fitness between hybrids and the S. cerevisiae control were tested with Z-tests (Bonferroni-adjusted P-values). Plots were generated with ggplot2.

### Synteny analysis and gene size variation

Synteny between *S. cerevisiae* S288C and *S. paradoxus* CBS432 was inferred with SynChro [44] using protein sequences; input FASTA/GFF files were pre-formatted with a custom Perl script (courtesy of L. Morales and I. Sedeño, LIIGH). From GFF annotations, we extracted CDS lengths for the ∼5,800 well-annotated *S. cerevisiae* genes and from *S. paradoxus* reference. We identified 5,366 one-to-one orthologs, 434 *S. cerevisiae*-only, and 112 *S. paradoxus*-only genes. For orthologs, absolute CDS length differences were computed and standardized (Z-scores across genes); outliers were defined as |Z| > 2 (≥2 SD from the mean).

### DNA extraction, library preparation, and sequencing

From each evolved population we isolated a single colony; genomic DNA from this clone was used for sequencing. Genomic DNA was extracted by phenol–chloroform, ethanol-precipitated, and treated with RNase A. Libraries were prepared as described previously [43]. Sequencing was performed by the Lewis-Sigler Institute for Integrative Genomics (Princeton University) core facility. Sequencing was done in three batches: (i) parental recombinant-hybrid progeny and their parental strains; (ii) polyploid evolved populations at generation 2,400; and (iii) haploid evolved populations at generation 5,000.

### Sequencing analysis pipeline

Whole-genome sequencing and analysis followed pipelines described previously for experimental evolution in *S. cerevisiae* [45] with minor adaptations for hybrid genomes. Illumina 150 bp paired-end reads were trimmed with Trimmomatic (v0.36) and aligned with BWA-MEM (v0.7.15) to a concatenated reference comprising the *S. cerevisiae* S288C and *S. paradoxus* CBS432 nuclear genomes plus both mitochondrial genomes (downloaded May 2022 from the Yeast PacBio resource: https://yjx1217.github.io/Yeast_PacBio_2016/data/). PCR duplicates were removed and per-site coverage was computed with samtools. Variants were called with FreeBayes (v1.1.0), filtered to remove low-quality calls, shared background variants (VCFtools v0.1.15; vcf-isec), and sites in low-complexity or recombination-breakpoint regions, and then annotated with SnpEff (v5.0). Zygosity was inferred from allele frequencies using a ≥0.9 threshold and a binomial test *(p < 0.001*).

For ancestry inference, gene-level coverage was extracted (samtools coverage), normalized, and the species of origin for each gene was assigned from the ratio of reads mapping to the *S. cerevisiae* versus *S. paradoxus* references (Supplementary Data 1); regions spanning recombination breakpoints that remained ambiguous were assigned to the species with higher mapped read support, assuming 2:2 segregation across contiguous gene blocks. Mitochondrial genotype was determined by mapping reads to the *S. cerevisiae* and *S. paradoxus* mitochondrial genomes at three conserved loci (*ATP6, COX2, COX1)* as described in Ref. [46].

### Hybrid complex composition and mutation enrichment analysis

We constructed a complex–gene map by integrating annotations from SGD, Complex Portal and published compendia [26, 47, 48] mapping systematic gene names and their associated protein complexes and allowing individual genes to participate in multiple complexes (Supplementary Data 3). Of 5,366 one-to-one orthologs, 2,121 genes mapped to 652 obligate complexes and 3,245 were classified as non-complex. Because complexes were defined strictly from one-to-one orthologs, subunits lacking an annotated *S. paradoxus* ortholog were excluded. As a result, 24 complexes differed in their apparent subunit number, and the final ortholog-based complex set comprised 628 complexes.

For each segregant, we read allele calls for all subunits and labeled the complex hybrid if it contained at least one *S. cerevisiae* and at least one *S. paradoxus* allele; otherwise, non-hybrid. This yielded a complex-by-strain hybrid-status matrix (Supplementary Data 3). To contextualize the observed hybridization patterns, we generated null expectations under random inheritance (independent per-gene segregation) and compared observed vs expected distributions of “hybrid background frequency” (number of segregants in which a complex is hybrid), stratified by complex size (2-subunits). To minimize linkage effects, analyses were repeated on unlinked complexes (all subunit genes on different chromosomes). We define essential complexes as ones where either protein is classified as essential by the *Saccharomyces* Genome Database ((yeastgenome.org).

For mutation enrichment, we compared the observed number of missense mutations in complexes classified as hybrid vs uniparental to expectations under a length-weighted null model and summarized effects as log_2_ fold-change (observed/expected), with corresponding log-transformed P-values. Complexes showing a significant excess of mutations when assembled in hybrid form were flagged as candidates for hybrid-specific adaptation.

### Identification of common targets

Candidate targets of selection were identified using the length-weighted Poisson as described previously [45]. Briefly, we tested each annotated *S. cerevisiae* CDS (∼5,800 genes) for enrichment of nonsynonymous mutations (missense, nonsense and frameshift) and, in a separate analysis, enrichment of missense-only mutations, with expected counts scaled by coding-sequence length (mean CDS length of the *S. cerevisiae/S. paradoxus* ortholog pair to approximate mutational opportunity). *S. paradoxus*–specific ORFs were excluded. Genes with adjusted *p < 10^-4^* and ≥10 nonsynonymous or missense mutations were classified as “common targets”. To distinguish hybrid-specific from general targets, we applied the same analysis to our evolved parental populations and to published *S. cerevisiae* evolution datasets [32, 34, 35] and compared gene-wise enrichment across datasets.

### Sequence alignment of candidate targets of selection

Protein sequences of candidate targets were aligned with MAFFT. Alignments (FASTA) were parsed with a custom Python script (Biopython) to compute per-gene statistics: percent identity, numbers of matches/mismatches/indels, total alignment length, and effective gene size. For position-wise divergence, alignments were reshaped to a residue-level table classifying each position as match, mismatch, or gap. Divergence tracks were visualized in R (ggplot2) using line/tile glyphs along the protein coordinate.

### LOF/non-LOF selection (Bayesian analysis)

We assessed whether selection in hybrids preferentially favored loss-of-function (LOF) versus non-LOF changes using a Bayesian model, which uses priors derived from a meta-analysis of laboratory evolution targets [45]. From our hybrid dataset, we took the 100 most recurrently mutated genes across segregants and classified coding mutations as LOF-consistent (nonsense, frameshift, start-loss, stop-gain) or non-LOF-consistent (missense); synonymous and intergenic variants were excluded.

### Variance scores analysis

To identify candidate genes under functional constraint or directional selection, we analyzed dispersion of nonsynonymous (missense-inclusive) mutation counts across all evolved hybrid populations. For each gene, we computed the mean, variance, and variance-to-mean ratio (VMR = variance/mean). Under a neutral Poisson model, strong overdispersion (VMR > 2) was flagged as indicative of population-specific adaptation or positive selection.

### Genome editing with CRISPR–Cas9

Targeted edits were introduced into parental and hybrid strains using pML104, which encodes constitutive Cas9 and a nourseothricin (*NatMX*) marker for selection. Species-specific single-guide RNAs (sgRNAs) were designed to target *AMN1, ADE1*, and *HSP104* and co-transformed with 500-bp donor DNA (gBlocks) carrying the desired substitutions. To prevent Cas9 re-cutting, donor templates included synonymous changes within the PAM or guide recognition sequence. Edited clones were selected on *NatMX* and verified by Sanger sequencing. For *HSP104* reconstructions, edits were recovered in the *S. cerevisiae* background; two *S. paradoxus*-targeting guides repeatedly failed to yield edited clones.

## DATA AVAILABILITY

The short-read sequencing data reported in this study have been deposited to the NCBI BioProject database. The parental genomes and the hybrid at generation 0 are in accession number PRJNA1330885. Sequencing of generation 2,400 of the polyploid hybrids is in accession number PRJNA1331785. Sequencing of generation 5,000 of the haploid hybrids is in accession number PRJNA1331855

## ACKNOWLEDGEMENTS AND FUNDING

We thank members of the Lang Lab for lab discussion and comments on the manuscript. This study was supported by a grant from the National Institutes of Health (R35GM149540). Portions of this research were conducted on Lehigh University’s Research Computing infrastructure partially supported by the National Science Foundation (Award 2019035).

## AUTHOR CONTRIBUTIONS

Conceptualization: GIL

Formal Analysis: AAM, GIL

Investigation: AAM, GIL

Writing – Original Draft Preparation: AAM, GIL

Writing – Review & Editing: AAM, GIL

Visualization: AAM, GIL

Supervision: GIL

Funding Acquisition: GIL

## COMPETING INTERESTS

The authors declare no competing interests.

## Notes

### Competing Interest Statement

The authors have declared no competing interest.

## REFERENCES

1. Maheshwari, S. and D.A. Barbash, The genetics of hybrid incompatibilities. Annual review of genetics, 2011. 45(1): p. 331–355.

2. Presgraves, D.C., The molecular evolutionary basis of species formation. Nature Reviews Genetics, 2010. 11(3): p. 175–180.

3. Abbott, R., et al., Hybridization and speciation. Journal of evolutionary biology, 2013. 26(2): p. 229–246.

4. Runemark, A., M. Vallejo-Marin, and J.I. Meier, Eukaryote hybrid genomes. PLoS genetics, 2019. 15(11): p. e1008404.

5. Servedio, M.R., J. Hermisson, and G.S. van Doorn, Hybridization may rarely promote speciation. Journal of Evolutionary Biology, 2013. 26(2): p. 282–285.

6. Bozdag, G.O. and J. Ono, Evolution and molecular bases of reproductive isolation. Current opinion in genetics & development, 2022. 76: p. 101952.

7. Maclean, C.J. and D. Greig, Prezygotic reproductive isolation between Saccharomyces cerevisiae and Saccharomyces paradoxus. BMC Evolutionary Biology, 2008. 8(1): p. 1.

8. Langdon, Q.K., et al., Fermentation innovation through complex hybridization of wild and domesticated yeasts. Nature Ecology & Evolution, 2019. 3(11): p. 1576–1586.

9. Molinet, J. and R. Stelkens, The evolution of thermal performance curves in response to rising temperatures across the model genus yeast. Proceedings of the National Academy of Sciences, 2025. 122(21): p. e2423262122.

10. Peris, D., et al., Hybridization and adaptive evolution of diverse Saccharomyces species for cellulosic biofuel production. Biotechnology for Biofuels, 2017. 10(1): p. 78.

11. Piatkowska, E.M., et al., Chimeric protein complexes in hybrid species generate novel phenotypes. PLoS Genetics, 2013. 9(10): p. e1003836.

12. D’Angiolo, M., et al., A yeast living ancestor reveals the origin of genomic introgressions. Nature, 2020. 587(7834): p. 420–425.

13. Heil, C.S.S., et al., Loss of heterozygosity drives adaptation in hybrid yeast. Molecular biology and evolution, 2017. 34(7): p. 1596.

14. Leducq, J.-B., et al., Speciation driven by hybridization and chromosomal plasticity in a wild yeast. Nature Microbiology, 2016. 1(1): p. 1–10.

15. Morales, L. and B. Dujon, Evolutionary role of interspecies hybridization and genetic exchanges in yeasts. Microbiology and Molecular Biology Reviews, 2012. 76(4): p. 721–739.

16. Chou, J.-Y., et al., Multiple molecular mechanisms cause reproductive isolation between three yeast species. PLoS biology, 2010. 8(7): p. e1000432.

17. Greig, D., A Screen for Recessive Speciation Genes Expressed in the Gametes of F1 Hybrid Yeast. PLOS Genetics, 2007. 3(2): p. e21.

18. Greig, D., et al., Hybrid speciation in experimental populations of yeast. Science, 2002. 298(5599): p. 1773–5.

19. Hou, J., et al., Comprehensive survey of condition-specific reproductive isolation reveals genetic incompatibility in yeast. Nature Communications, 2015. 6(1): p. 7214.

20. Kao, K.C., K. Schwartz, and G. Sherlock, A genome-wide analysis reveals no nuclear Dobzhansky-Muller pairs of determinants of speciation between S. cerevisiae and S. paradoxus, but suggests more complex incompatibilities. PLoS genetics, 2010. 6(7): p. e1001038.

21. Lee, H.-Y., et al., Incompatibility of nuclear and mitochondrial genomes causes hybrid sterility between two yeast species. Cell, 2008. 135(6): p. 1065–1073.

22. Delneri, D., et al., Engineering evolution to study speciation in yeasts. Nature, 2003. 422(6927): p. 68–72.

23. Fischer, G., et al., Highly Variable Rates of Genome Rearrangements between Hemiascomycetous Yeast Lineages. PLOS Genetics, 2006. 2(3): p. e32.

24. Hunter, N., et al., The mismatch repair system contributes to meiotic sterility in an interspecific yeast hybrid. The EMBO journal, 1996. 15(7): p. 1726–1733.

25. Bozdag, G.O., et al., Breaking a species barrier by enabling hybrid recombination. Current biology, 2021. 31(4): p. R180–R181.

26. Swamy, K.B., S.C. Schuyler, and J.-Y. Leu, Protein complexes form a basis for complex hybrid incompatibility. Frontiers in genetics, 2021. 12: p. 609766.

27. Tang, S. and D.C. Presgraves, Evolution of the Drosophila nuclear pore complex results in multiple hybrid incompatibilities. Science, 2009. 323(5915): p. 779–782.

28. Tirosh, I., et al., A Yeast Hybrid Provides Insight into the Evolution of Gene Expression Regulation. Science, 2009. 324(5927): p. 659–662.

29. Brion, C., et al., Variation of the meiotic recombination landscape and properties over a broad evolutionary distance in yeasts. PLoS Genet, 2017. 13(8): p. e1006917.

30. Fang, O., et al., Amn1 governs post-mitotic cell separation in Saccharomyces cerevisiae. PLoS genetics, 2018. 14(10): p. e1007691.

31. Gonçalves, P., et al., Evidence for divergent evolution of growth temperature preference in sympatric Saccharomyces species. PloS one, 2011. 6(6): p. e20739.

32. Fisher, K.J., et al., Adaptive genome duplication affects patterns of molecular evolution in Saccharomyces cerevisiae. PLoS genetics, 2018. 14(5): p. e1007396.

33. Harari, Y., et al., Spontaneous changes in ploidy are common in yeast. Current Biology, 2018. 28(6): p. 825–835. e4.

34. Johnson, M.S., et al., Phenotypic and molecular evolution across 10,000 generations in laboratory budding yeast populations. Elife, 2021. 10: p. e63910.

35. Marad, D.A., S.W. Buskirk, and G.I. Lang, Altered access to beneficial mutations slows adaptation and biases fixed mutations in diploids. Nature Ecology & Evolution, 2018. 2(5): p. 882–889.

36. Dandage, R., et al., Frequent assembly of chimeric complexes in the protein interaction network of an interspecies yeast hybrid. Molecular biology and evolution, 2021. 38(4): p. 1384–1401.

37. Li, C., Z. Wang, and J. Zhang, Toward genome-wide identification of Bateson-Dobzhansky-Muller incompatibilities in yeast: a simulation study. Genome Biol Evol, 2013. 5(7): p. 1261–72.

38. Xu, M. and X. He, Genetic incompatibility dampens hybrid fertility more than hybrid viability: yeast as a case study. PLoS One, 2011. 6(4): p. e18341.

39. Jhuang, H.Y., H.Y. Lee, and J.Y. Leu, Mitochondrial-nuclear co-evolution leads to hybrid incompatibility through pentatricopeptide repeat proteins. EMBO Rep, 2017. 18(1): p. 87–101.

40. Kowalec, P., et al., Newly identified protein Imi1 affects mitochondrial integrity and glutathione homeostasis in Saccharomyces cerevisiae. FEMS Yeast Research, 2015. 15(6): p. fov048.

41. Liti, G., et al., Segregating YKU80 and TLC1 Alleles Underlying Natural Variation in Telomere Properties in Wild Yeast. PLOS Genetics, 2009. 5(9): p. e1000659.

42. Sawyer, S.L. and H.S. Malik, Positive selection of yeast nonhomologous end-joining genes and a retrotransposon conflict hypothesis. Proceedings of the National Academy of Sciences, 2006. 103(47): p. 17614–17619.

43. Buskirk, S.W., R.E. Peace, and G.I. Lang, Hitchhiking and epistasis give rise to cohort dynamics in adapting populations. Proc Natl Acad Sci U S A, 2017. 114(31): p. 8330–8335.

44. Drillon, G., A. Carbone, and G. Fischer, SynChro: A Fast and Easy Tool to Reconstruct and Visualize Synteny Blocks along Eukaryotic Chromosomes. PLOS ONE, 2014. 9(3): p. e92621.

45. Martínez, A.A. and G.I. Lang, Identifying Targets of Selection in Laboratory Evolution Experiments. Journal of Molecular Evolution, 2023. 91(3): p. 345–355.

46. Yue, J.X., et al., Contrasting evolutionary genome dynamics between domesticated and wild yeasts. Nat Genet, 2017. 49(6): p. 913–924.

47. Laurent, J.M., et al., Humanization of yeast genes with multiple human orthologs reveals functional divergence between paralogs. PLOS Biology, 2020. 18(5): p. e3000627.

48. Swamy, K.B., et al., Proteotoxicity caused by perturbed protein complexes underlies hybrid incompatibility in yeast. Nature Communications, 2022. 13(1): p. 4394.

