## Supplementary Figure 1 for "Weak pervasive incompatibilities and compensatory adaptation drive hybrid genome evolution in yeast"

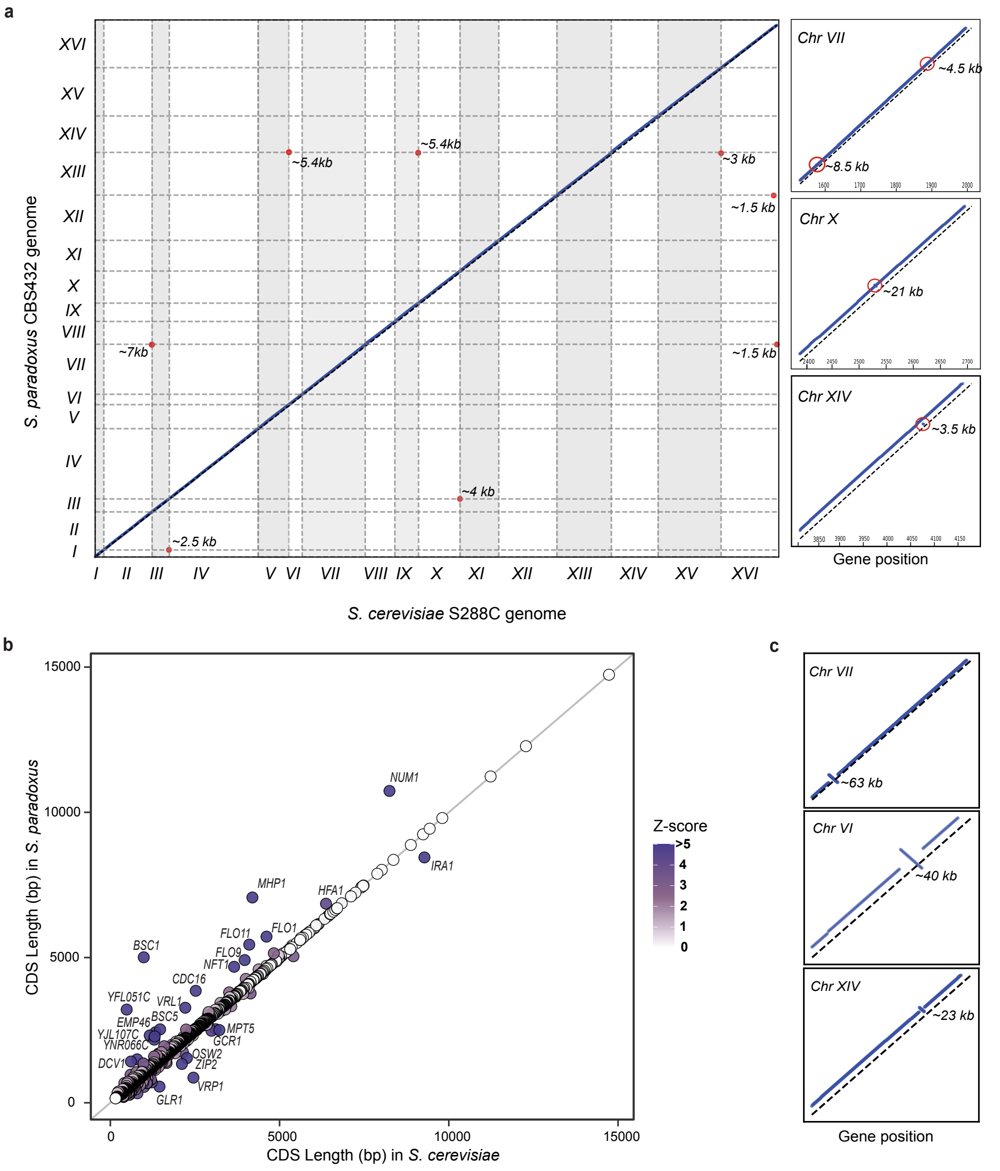


**Supplementary Fig 1 | Synteny conservation between *S. cerevisiae* S288C and *S. paradoxus* CBS432. a,** Synteny blocks between S288C and CBS432. Red dots indicate the positions and approximate sizes of chromosomal translocations. An inset of chromosomes VII, X and XIV highlights regions containing inversions. **b,** Comparison of gene lengths for 5,366 one-to-one orthologous genes between the two species. Z-scores of length differences were calculated to assess significant size variation; 57 genes showed absolute Z-scores > 2, indicating substantial size differences. Genes with absolute Z-scores > 5 are labeled. **c,** Zoomed views of chromosomes XIV, VII and VI illustrate inversions in *S. paradoxus* YPS138, shown for comparison with the *S. paradoxus* reference strain CBS432 used in this study.
