## Supplementary Figure 2 for "Weak pervasive incompatibilities and compensatory adaptation drive hybrid genome evolution in yeast"

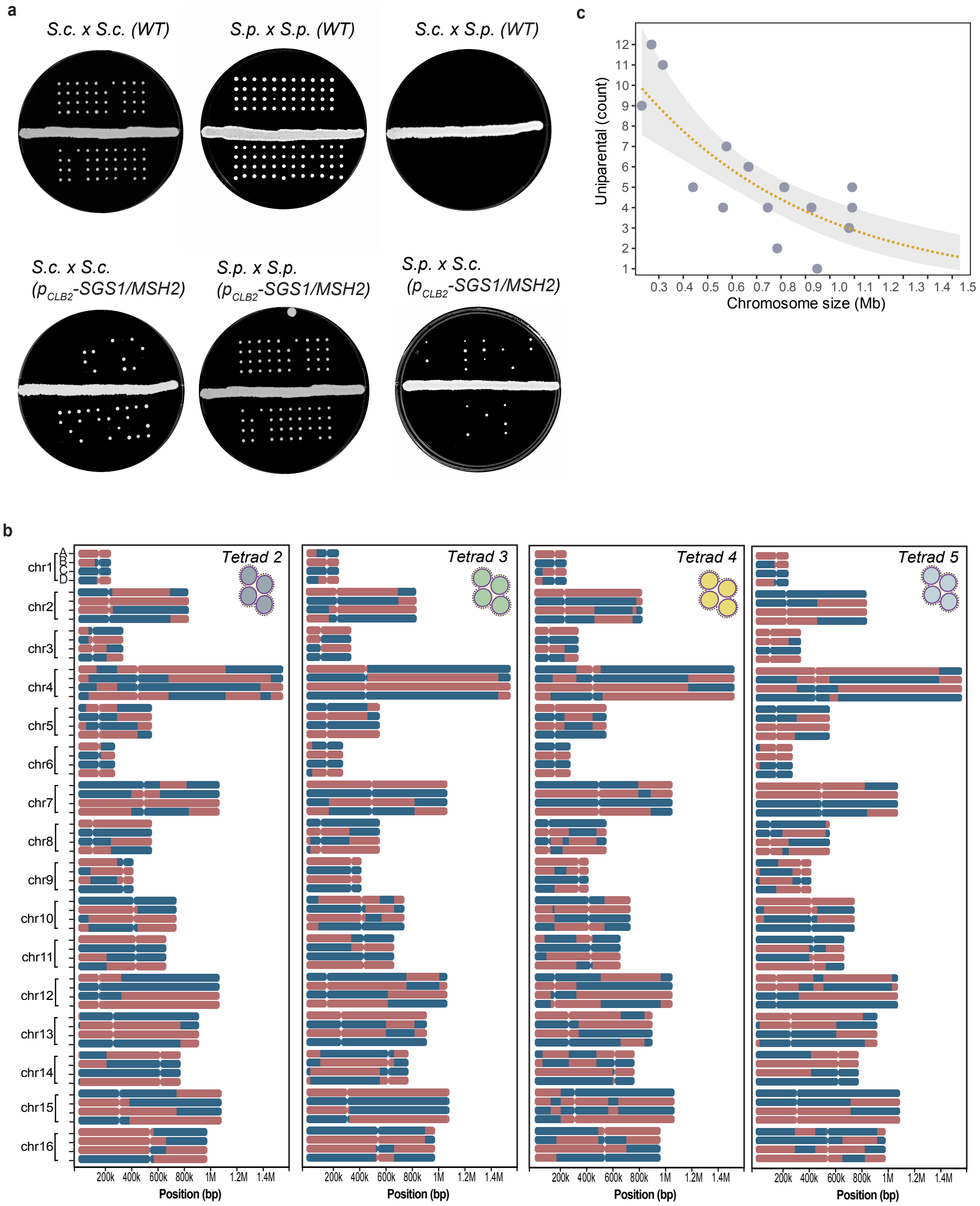


**Supplementary Fig 2 | Tetrad dissection and hybrid genome architecture. a,** Tetrad dissection of intra- and interspecific crosses between *S. paradoxus* CSB432 (*S.p.*) and *S. cerevisiae* S288C (*S.c.*) with and without suppression of *MSH2* and *SGS1*. **b,** Collapsed coverage tracks summarizing genomic blocks and recombination maps for Tetrads 2–5. Blue segments indicate *S. cerevisiae* inheritance; pink segments indicate *S. paradoxus* inheritance, highlighting crossover tracts and uniparental chromosomal blocks. **c,** Relationship between chromosome size and uniparental events. Each point represents a chromosome (x-axis: size in Mb; y-axis: number of uniparental occurrences across the 20 segregants). The solid line shows a Poisson generalized linear fit (log link) with a shaded 95% confidence interval.The negative slope indicates that uniparental events are more frequent on smaller chromosomes and decline with increasing chromosome size.
