## Supplementary Figure 3 for "Weak pervasive incompatibilities and compensatory adaptation drive hybrid genome evolution in yeast"

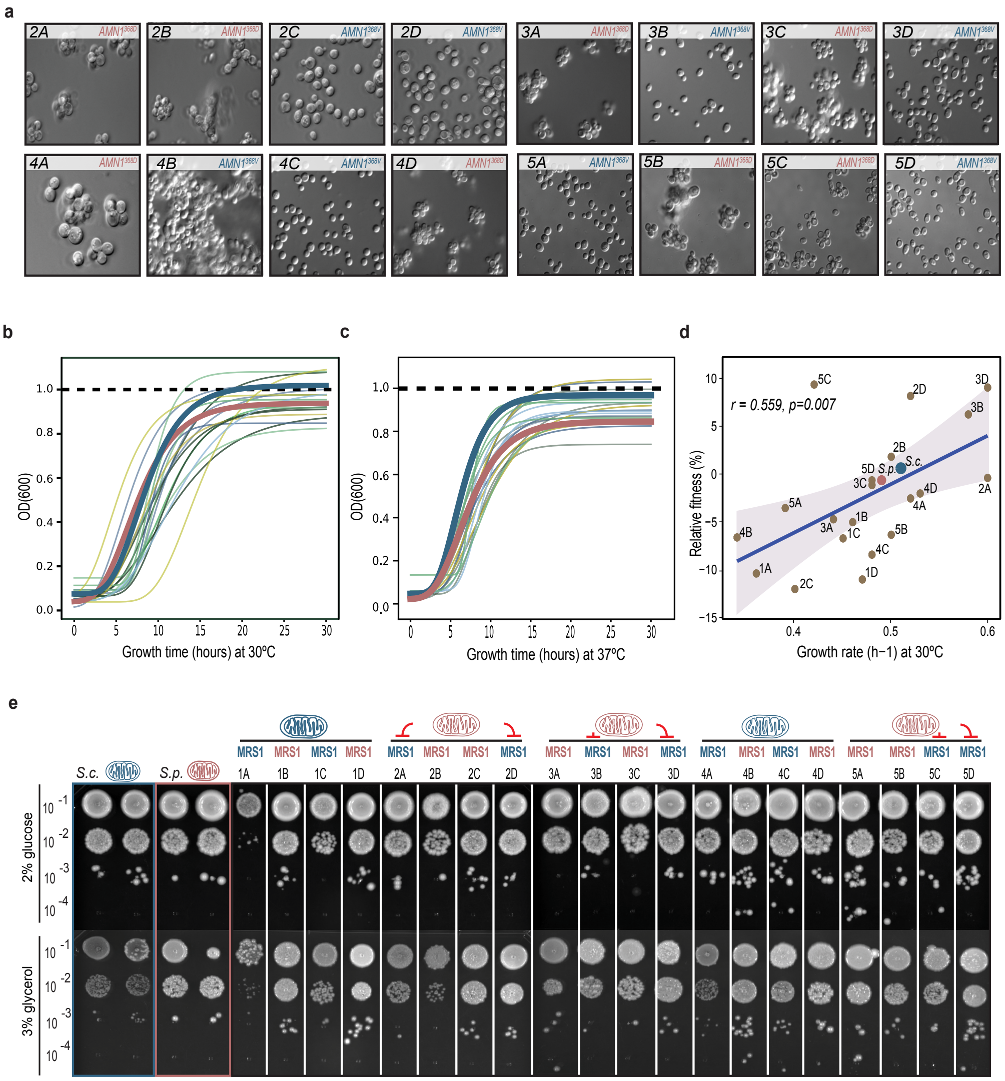


**Supplementary Fig 3 | Phenotypic segregation and mitochondrial ancestry in haploid segregants. a,** Microscopy of segregants from tetrads 2–5 shows 2:2 segregation of the clumpy phenotype, consistent with *AMN1* allele inheritance identified by whole-genome sequencing as a candidate for mother–daughter separation defects.The *S. paradoxus AMN1$^{368D}$* allele perfectly tracks with clumping, whereas the *S. cerevisiae AMN1$^{368V}$* allele yields normal separation. The degree of clumpiness varies among segregants, indicating additional genetic modifiers and/or background effects. Segregant 4B forms large clumps despite lacking the *S. paradoxus* allele, suggesting an alternative genetic cause in that background. **b,** Growth curves at 30 °C and **c,** 37 °C in rich media for 20 segregants from five tetrads; colours distinguish individual segregants. **d,** Relationship between competitive fitness and growth rate at 30 °C (Pearson's r = 0.559, *p = 0.007*), indicating a moderate positive correlation. **e,** Spot assays on glucose (YPD) and glycerol (YP + 3% glycerol) at 30 °C. Mitochondrial origin is colour-coded: *S. cerevisiae* (blue), *S. paradoxus* (pink). All segregants from a given tetrad share the same uniparentally inherited mitochondrial genome. The inherited allele of *MRS1* is shown for each segregant; the reported incompatibility arises when *S. paradoxus* mitochondria co-occur with the *S. cerevisiae MRS1* allele. Despite the presence of this combination in several segregants, we did not observe clear mitochondrial–nuclear incompatibility under these conditions.
