## Supplementary Figure 4 for "Weak pervasive incompatibilities and compensatory adaptation drive hybrid genome evolution in yeast"

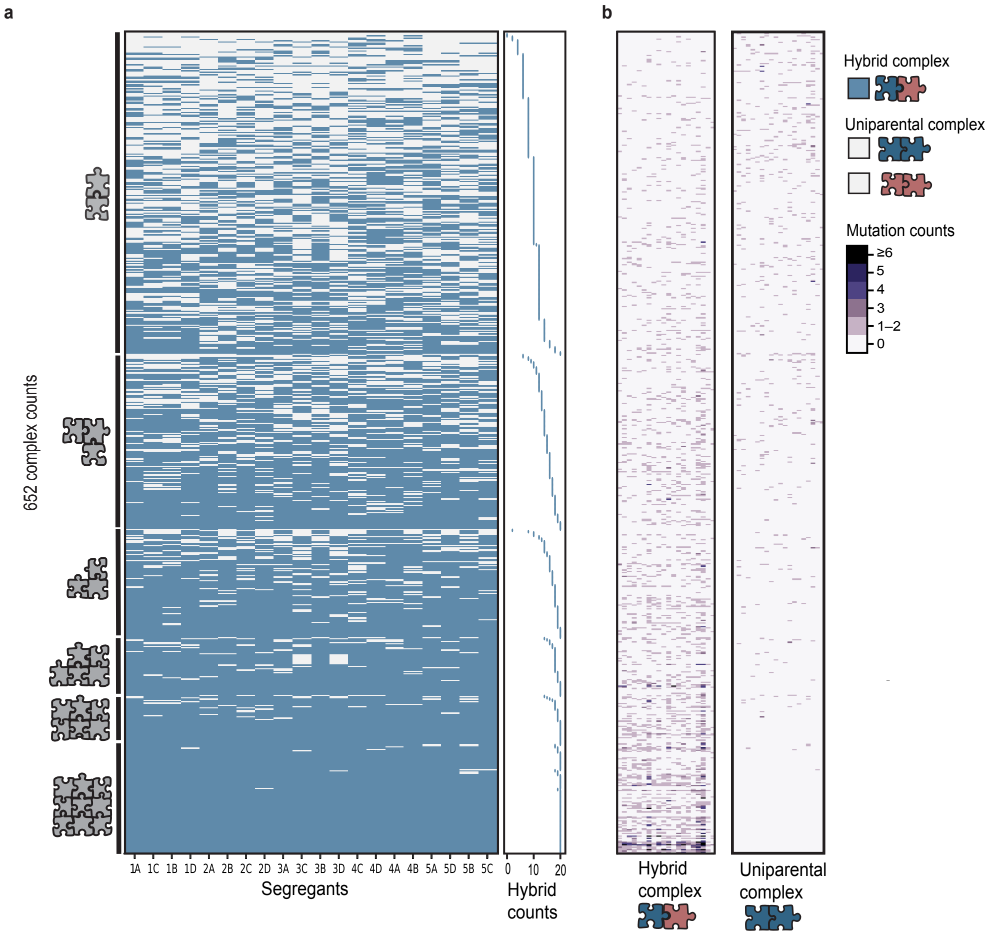


**Supplementary Fig. 4 | Hybrid protein-complex composition and mutational targeting in haploid segregants. a,** Hybrid versus uniparental composition of 626 conserved protein complexes across 20 recombinant segregants. Complexes are ordered by subunit number and, within each size class, by the number of segregants in which the complex is hybrid. Blue cells indicate complexes that assemble with subunits from both *S. cerevisiae* and *S. paradoxus* in a given segregant (hybrid complexes), whereas white cells indicate complexes composed entirely of subunits from a single species (uniparental complexes). The adjacent strip shows, for each complex, the total number of segregants (0–20) in which it forms a hybrid complex. Smaller assemblies (2–4 subunits) show much greater variability in hybrid status than larger assemblies: across 13,040 complex–segregant combinations, 2–4-subunit complexes are hybrid in ~62% of cases, whereas complexes with ≥5 subunits are hybrid in ~96%. A logistic regression of hybrid status on subunit number confirms this strong size dependence (odds ratio ≈ 2.27 per additional subunit, 95% CI 2.17–2.37, $p < 10^{-16}$). **b,** Heat map of missense mutations mapped to hybrid and uniparental complexes in evolved populations. Darker purple shading denotes complexes that accumulate more missense mutations across evolved populations.
