## Supplementary Figure 5 for "Weak pervasive incompatibilities and compensatory adaptation drive hybrid genome evolution in yeast"

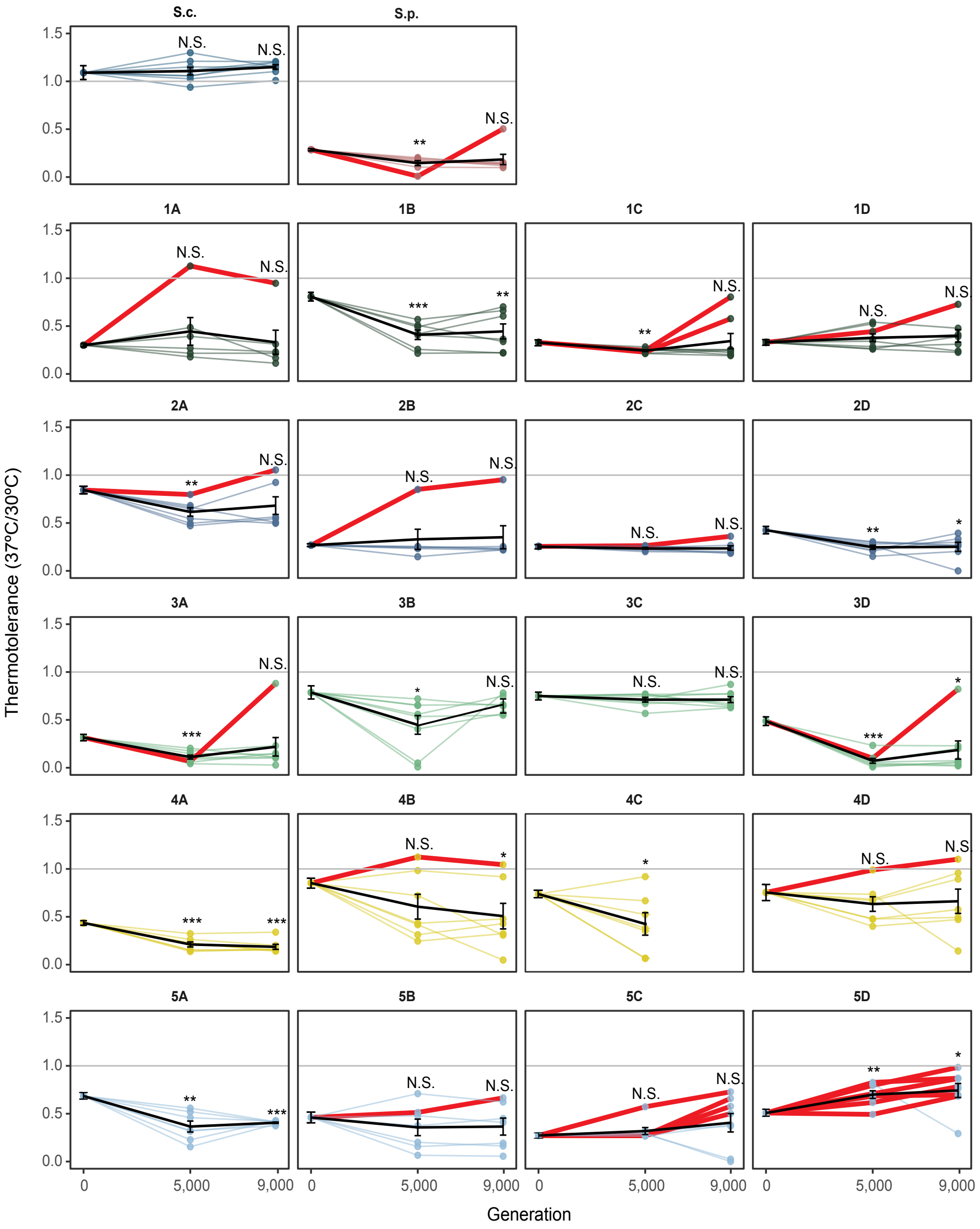


**Supplementary Fig. 5 | Thermotolerance trajectories of evolved populations.** Thermotolerance (R=S37/S30) was calculated from spot assays as the mean growth at 37 °C relative to 30 °C across four 1:32 serial dilutions. For each founder, all generation-0 replicates were collapsed to a single ancestral mean, and trajectories of independent populations are shown at generations 5,000 and 9,000. For each hybrid background, we calculated the mean thermotolerance (± s.d.) at generations 5,000 and 9,000 and performed paired t-tests to test whether thermotolerance at each generation differed significantly from the ancestral value (generation 0) across replicate lineages (*p < 0.05; *p < 0.01; **p < 0.001*; N.S., Not Significant). “Strong improver” lineages were defined as those whose final thermotolerance at generation 9,000 exceeded their hybrid’s ancestral mean by more than three ancestral standard deviations and are highlighted in red.
