## Supplementary Figure 6 for "Weak pervasive incompatibilities and compensatory adaptation drive hybrid genome evolution in yeast"

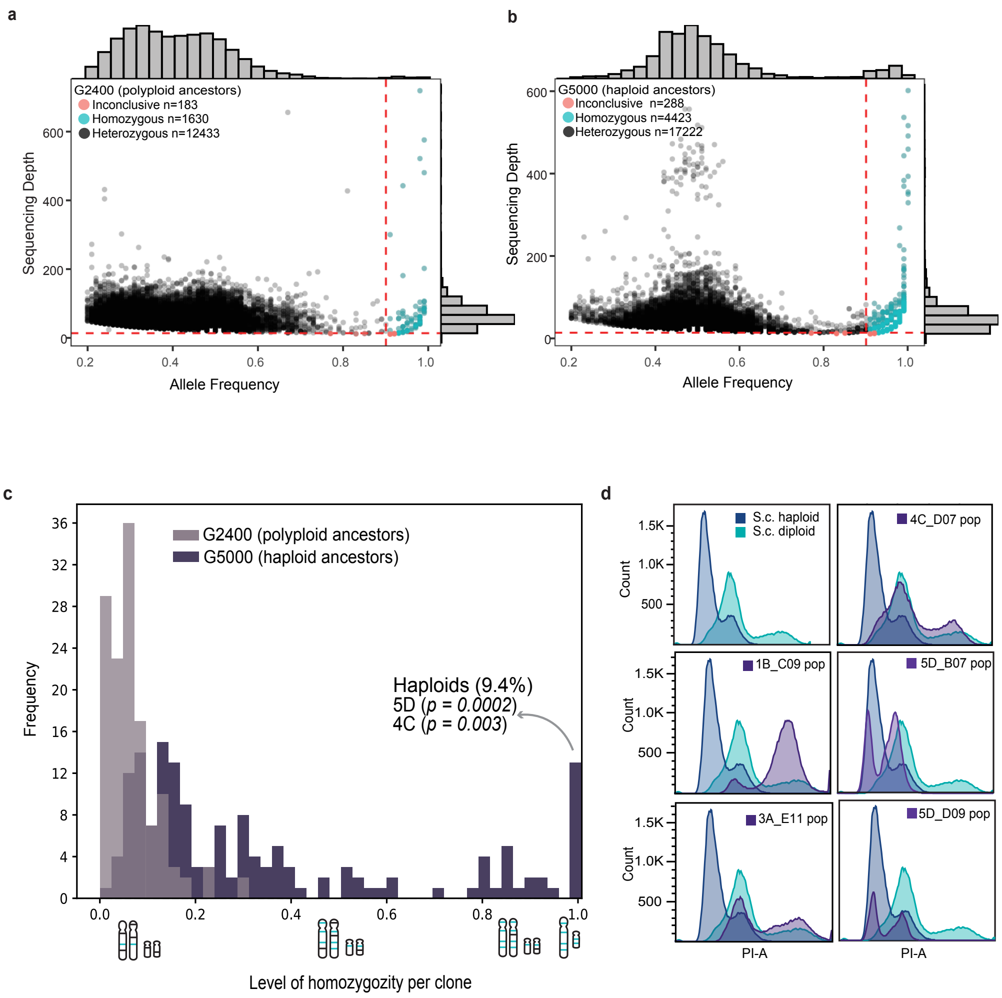


**Supplementary Fig. 6 | Evidence of auto-diploidization in evolved haploid populations. a,** Allele frequency and sequencing depth for 143 populations founded from segregant-derived diploids/polyploids and sequenced at generation 2,400. The x-axis shows allele frequency and the y-axis shows read depth for each called variant. The red dotted line indicates the threshold for classifying homozygous mutations ($p < 0.001$). Homozygous variants account for 11.44% of all mutations. Variants with normalized sequencing depth below 0.9 were classified as inconclusive. **b,** Equivalent analysis for 132 populations founded from haploid segregants and sequenced at generation 5,000. Homozygous mutations account for 20.17% of all mutations. **c,** Distribution of populations by mutation homozygosity across both datasets. The x-axis shows the fraction of mutations classified as homozygous in each population. Light purple, populations founded from diploid/polyploid ancestors (generation 2,400); dark purple, populations founded from haploid ancestors (generation 5,000). Haploid-founded populations show a marked shift toward higher fractions of homozygous mutations. Many populations retain substantial heterozygosity, consistent with autodiploidization occurring at different stages of the experiment. Overall, 90.6% of haploid-founded populations became autodiploid, whereas 9.4% remained haploid. Haploid retention is significantly enriched among replicates derived from ancestors 5D ($p = 0.0002$) and 4C ($p = 0.003$; Fisher's exact test). **d,** Flow-cytometry profiles of representative clones with high homozygosity confirm diploidization: most haploid-derived populations underwent autodiploidization, whereas a minority remained haploid (for example, background 5D).
