## Supplementary Figure 7 for "Weak pervasive incompatibilities and compensatory adaptation drive hybrid genome evolution in yeast"

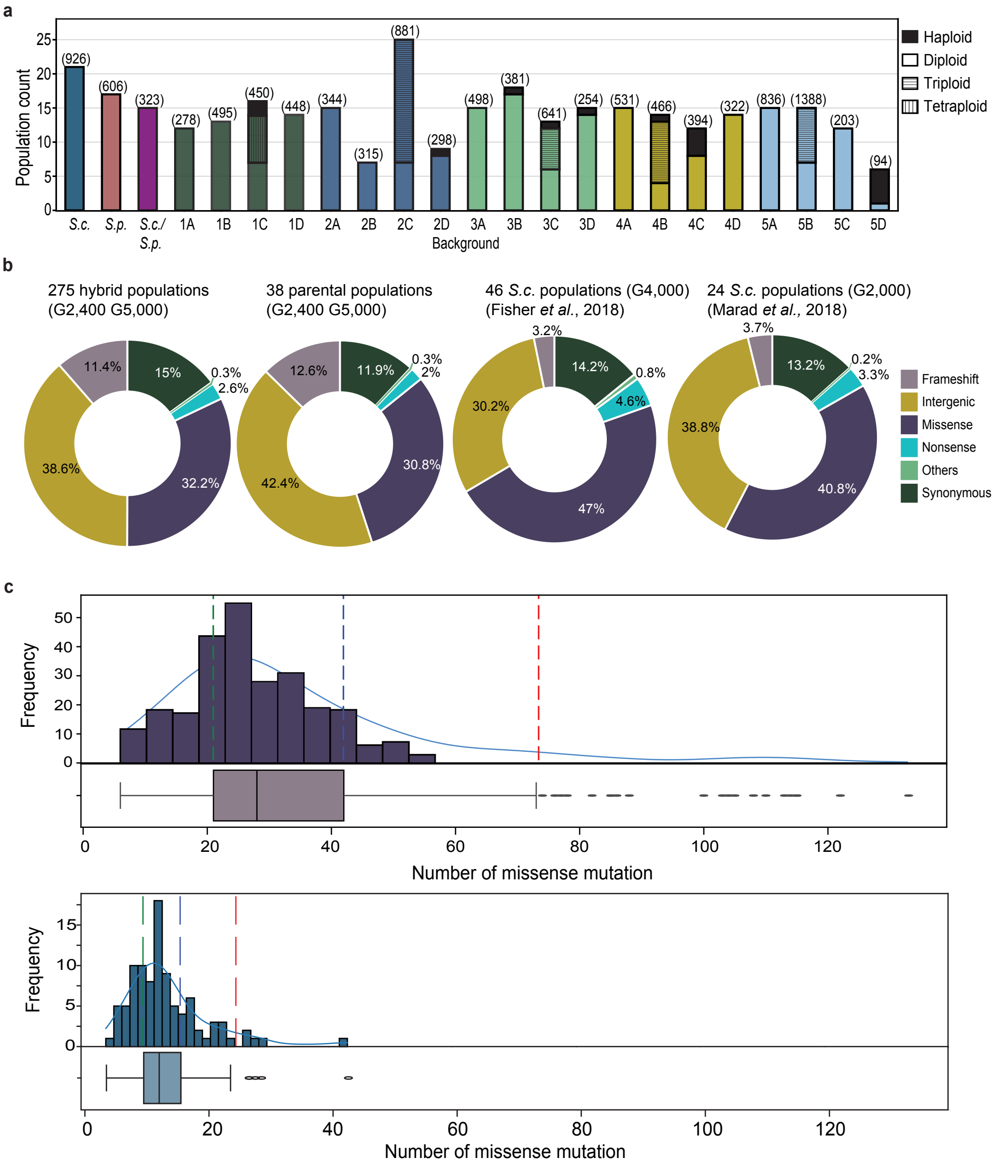


**Supplementary Fig. 7 | Evolved population counts and mutation spectra. a,** Number of evolved populations per genetic background. Black bars denote haploid-founded populations; vertical hatching marks triploid lines (derived from segregants 2C, 3C, 4B and 5B); horizontal hatching marks tetraploid lines (derived from segregant 1C). **b,** Mutation-class proportions across hybrid populations, parental controls and published datasets (Fisher et al., 2018: YPD autodiploids, n = 92; Marad et al., 2018: YPD haploids, n = 48). Our hybrids show a higher fraction of frameshifts than prior studies, consistent with a mutator state potentially arising from the temporal suppression of *SGS1/MSH2*. **c,** Distribution of missense mutations per evolved hybrid population (histogram with accompanying box plot). Mean ± s.d. = 34 ± 22. Outliers (Q3 + 1.5 × IQR) are flagged as putative mutator backgrounds; background 5B is significantly enriched relative to most others (two-sided Mann–Whitney U test with Bonferroni correction, adjusted $p < 0.001$), with 3C also enriched to a lesser extent. **d,** Distribution of total mutations per evolved clone in Fisher et al. (2018) for comparison (YPD autodiploids, n = 92); mean ± s.d. = 14 ± 6.
