## Supplementary Figure 8 for "Weak pervasive incompatibilities and compensatory adaptation drive hybrid genome evolution in yeast"

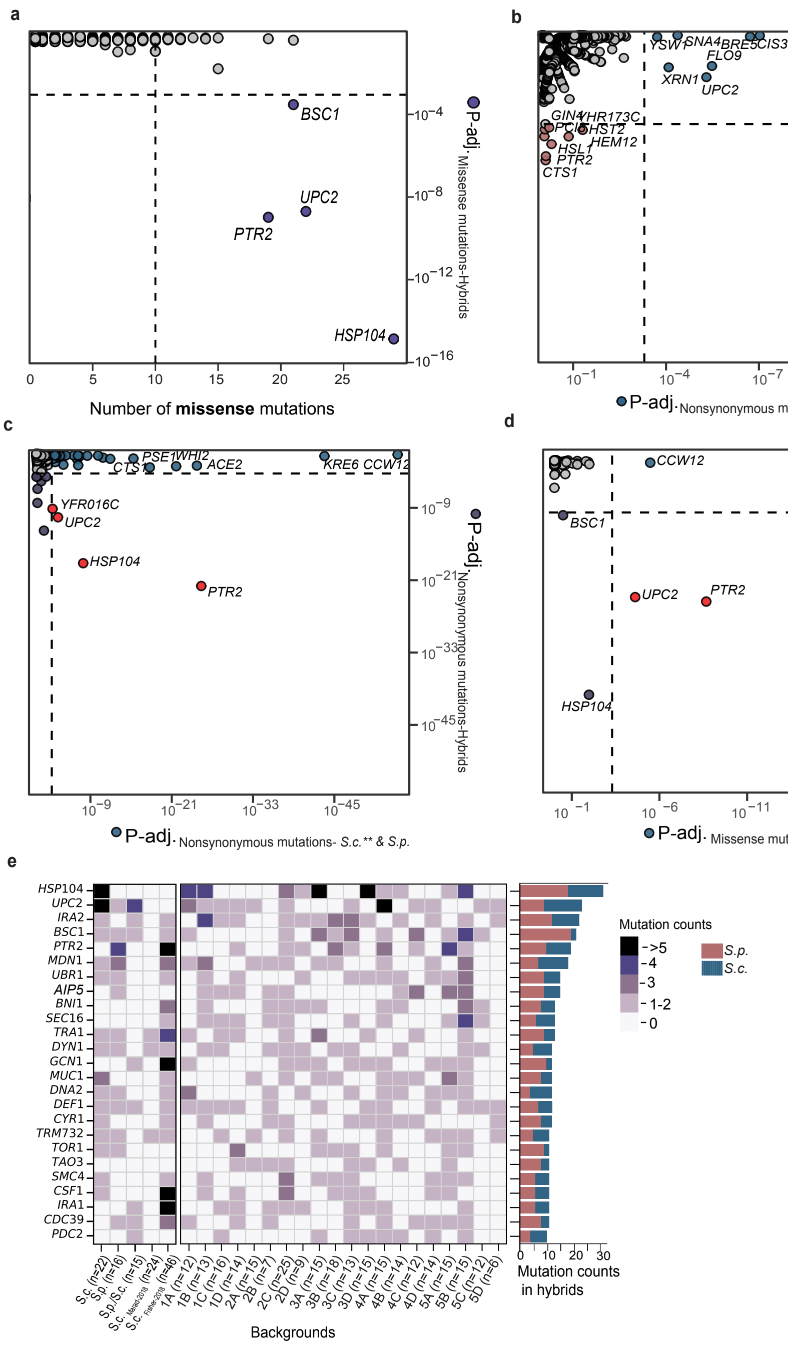


**Supplementary Fig. 8 | Targets of selection in evolved hybrid populations. a,** Genome-wide missense-enrichment analysis across ~5,800 coding sequences (CDSs) in hybrid populations under a Poisson model weighted by mean CDS length across species. The x-axis shows the observed missense count per gene; the y-axis shows the corresponding null probability. The dotted line marks the Benjamini–Hochberg FDR threshold ($p < 10^{-4}$); genes passing this threshold are highlighted in purple. **b,** Nonsynonymous enrichment in evolved parental populations (*S. cerevisiae*, blue; *S. paradoxus,* pink). Colour indicates the species in which each gene is significantly enriched for nonsynonymous mutations. **c,** Comparison of adjusted probabilities for nonsynonymous mutation enrichment between hybrids and parental populations, including published S. cerevisiae datasets (Fisher et al., 2018; Marad et al., 2018; Johnson et al., 2021). **d,** As in c, but restricted to missense mutations. **e,** Heat map of the 25 most frequently missense-mutated genes across hybrids, parental controls and reference datasets (Fisher et al., 2018; Marad et al., 2018). Bars at right show per-gene missense totals partitioned by allele of origin.
