## Supplementary Figure 9 for "Weak pervasive incompatibilities and compensatory adaptation drive hybrid genome evolution in yeast"

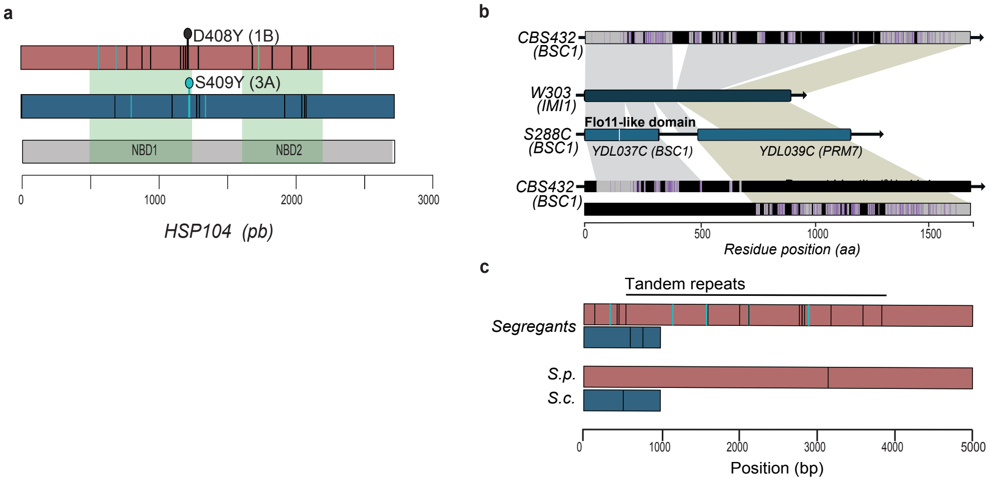


**Supplementary Fig. 9 | Mutational hotspots and background-specific selection. a,** *HSP104*: schematic of evolved mutations across parental and recombinant hybrid populations; heterozygous variants are shown as black ticks and homozygous variants as light blue ticks. **b,** *BSC1/PRM7/IMI1* architecture: CDS diagrams for *S. cerevisiae* (*BSC1* and *PRM7* in S288C; fused *IMI1* in W303) and the corresponding *S. paradoxus BSC1* gene. Amino-acid matches are shown in grey, mismatches in purple and gaps or regions absent in *S. cerevisiae* in black. The *S. paradoxus* gene aligns more closely to *IMI1* than to *BSC1/PRM7*, with low overall identity (~41%) and extensive tandem repeats. **c,** *BSC1*: schematic of evolved mutations across parental and segregant-founded populations; heterozygous variants are shown as black ticks and homozygous variants as light blue ticks.
